# A spatially mapped gene expression signature for intestinal stem-like cells identifies high-risk precursors of gastric cancer

**DOI:** 10.1101/2023.09.20.558462

**Authors:** Robert J. Huang, Ignacio A. Wichmann, Andrew Su, Anuja Sathe, Miranda V. Shum, Susan M. Grimes, Rithika Meka, Alison Almeda, Xiangqi Bai, Jeanne Shen, Quan Nguyen, Manuel R. Amieva, Joo Ha Hwang, Hanlee P. Ji

## Abstract

**Objective:** Gastric intestinal metaplasia (**GIM**) is a precancerous lesion that increases gastric cancer (**GC**) risk. The Operative Link on GIM (**OLGIM**) is a combined clinical-histopathologic system to risk-stratify patients with GIM. The identification of molecular biomarkers that are indicators for advanced OLGIM lesions may improve cancer prevention efforts.

**Methods:** This study was based on clinical and genomic data from four cohorts: 1) GAPS, a GIM cohort with detailed OLGIM severity scoring (N=303 samples); 2) the Cancer Genome Atlas (N=198); 3) a collation of in-house and publicly available scRNA-seq data (N=40), and 4) a spatial validation cohort (N=5) consisting of annotated histology slides of patients with either GC or advanced GIM. We used a multi-omics pipeline to identify, validate and sequentially parse a highly-refined signature of 26 genes which characterize high-risk GIM.

**Results:** Using standard RNA-seq, we analyzed two separate, non-overlapping discovery (N=88) and validation (N=215) sets of GIM. In the discovery phase, we identified 105 upregulated genes specific for high-risk GIM (defined as OLGIM III-IV), of which 100 genes were independently confirmed in the validation set. Spatial transcriptomic profiling revealed 36 of these 100 genes to be expressed in metaplastic foci in GIM. Comparison with bulk GC sequencing data revealed 26 of these genes to be expressed in intestinal-type GC. Single-cell profiling resolved the 26-gene signature to both mature intestinal lineages (goblet cells, enterocytes) and immature intestinal lineages (stem-like cells). A subset of these genes was further validated using single-molecule multiplex fluorescence *in situ* hybridization. We found certain genes (*TFF3* and *ANPEP*) to mark differentiated intestinal lineages, whereas others (*OLFM4* and *CPS1*) localized to immature cells in the isthmic/crypt region of metaplastic glands, consistent with the findings from scRNAseq analysis.

**Conclusions:** using an integrated multi-omics approach, we identified a novel 26-gene expression signature for high-OLGIM precursors at increased risk for GC. We found this signature localizes to aberrant intestinal stem-like cells within the metaplastic microenvironment. These findings hold important translational significance for future prevention and early detection efforts.

## INTRODUCTION

Gastric cancer (**GC**) is a leading source of global cancer morbidity and mortality.^1^ Survival from GC both worldwide and in Western nations remains poor (under 35%),^2^ largely due to advanced stages at time of diagnosis. The intestinal subtype makes up a substantial majority of GCs and follows a carcinogenic pathway termed Correa’s cascade.^3^ This premalignant evolution involves the gastric mucosa progressing through a series of histopathologic changes: non-atrophic gastritis (**NAG**), chronic atrophic gastritis (**CAG**), gastric intestinal metaplasia (**GIM**), dysplasia and ultimately GC.

GIM provides an opportunity for cancer interception, as it often persists and poses a continued risk for GC.^4–6^ The prevalence of GIM is estimated to be 5-10% of the general population in Western nations.^7,8^ However, only a subset of patients with GIM will progress to GC over long-term follow-up.^9–13^ Identifying this subset of “high-risk” GIM has become a high clinical priority and may lead to strategies of early detection and mortality reduction. One promising risk stratification tool is the Operative Link on Gastric Intestinal Metaplasia (**OLGIM**) staging framework. This validated scoring system incorporates both histologic severity and anatomic extent of GIM, with higher scores associated with higher subsequent cancer risk in both case-control^14,15^ and cohort studies.^16–18^ Citing an example, it was determined that advanced OLGIM lesions (Stages III or IV) are 25-fold more likely to progress to GC compared to early OLGIM lesions (Stage I).^18^ Understanding the biological underpinnings of high-risk GIM provides insights into the cellular mechanisms underlying the progression of GC and will yield high-value translational biomarkers for risk assessment.

*Helicobacter pylori* (**Hp**) is the etiologic agent most closely associated with intestinal-type GC,^19,20^ and most prior studies of gastric precursors have either focused on Hp-infected individuals or have drawn from populations with high Hp prevalence.^21–24^ However, Hp infection has been rapidly declining in Western nations, and most patients diagnosed with GIM have undergo bacterial eradication per clinical guidelines.^25^ Moreover, patients with GIM remain at significantly elevated risk even following Hp eradication.^5,6^ In this study, we developed a gene signature in patients without active Hp infection reflecting what is observed in western nations. This removed potential confounding variable in the relationship between advanced histology and transcriptomic profiling.

Using an OLGIM-staged cohort, we identified a gene expression signature which characterizes advanced, high-risk GIM lesions (defined as OLGIM III or IV) originating from Hp-negative individuals. This analysis used an integrated multi-omics approach that included conventional RNA sequencing (**RNA-seq**), spatial transcriptomics analysis, single-cell RNA sequencing (**scRNA-seq**), and single-molecule fluorescent *in situ* hybridization (**smFISH**). We orthogonally validated the RNAseq expression profiles of high-risk GIM to generate a highly-refined and spatially-resolved gene expression signature. Using scRNA-seq, these genes were mapped to specific cell populations in GIM lesions, including mature and stem-like intestinal lineages. A subset of these genes was further validated by smFISH. Overall, we discovered a spatially mapped high-risk gene expression signature which characterizes advanced, high-risk GIM lesions, is shared by intestinal-type GCs, and localizes to aberrant intestinal stem-like cells within the metaplastic microenvironment.

## MATERIALS AND METHODS

We provide an overview of both the experimental methods and the clinical cohorts used in this study in **Figure 1**. Complete methodologic details are provided in the **Supplementary Methods**.

**Figure 1:**
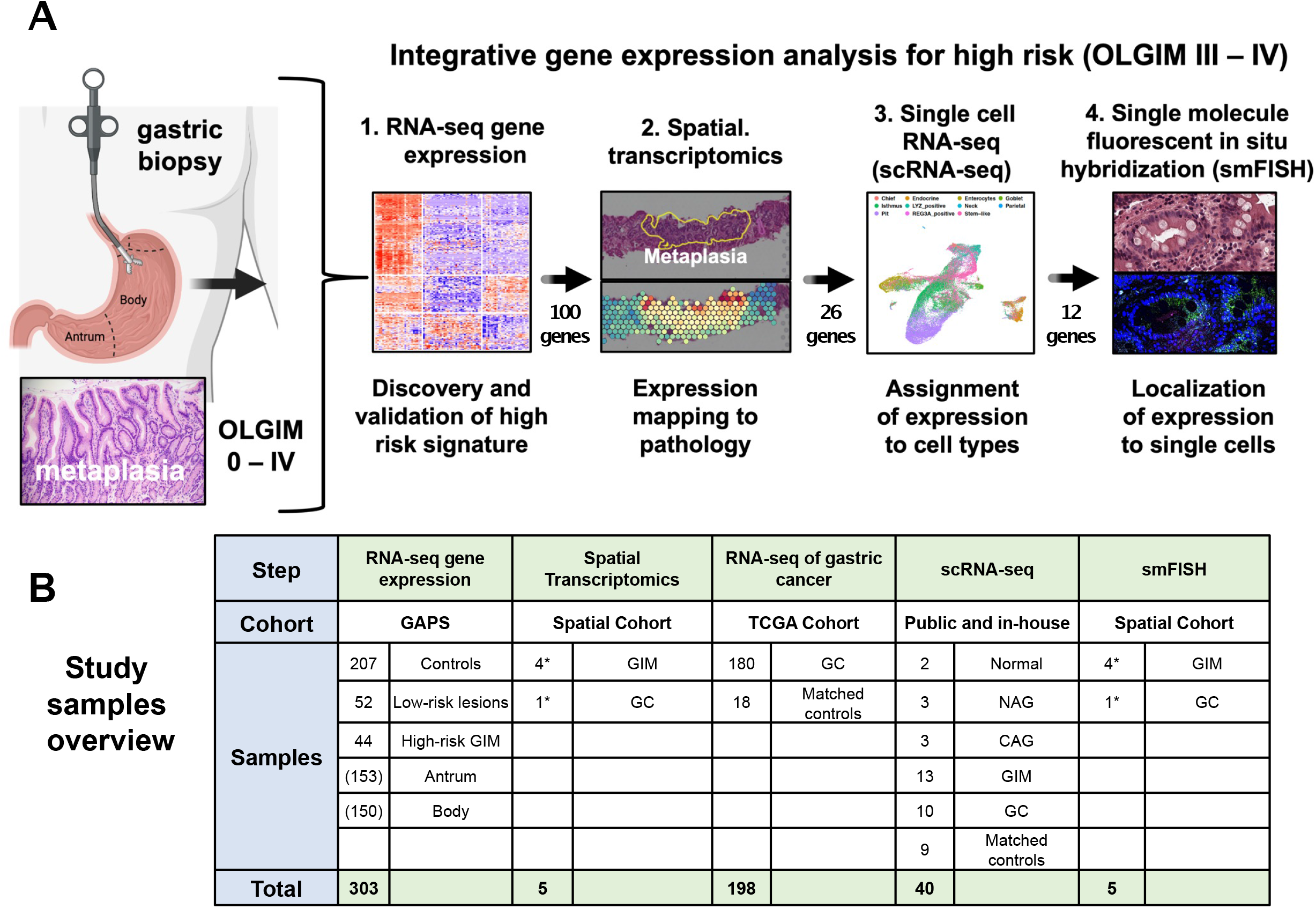
Overview of study design. **A)** Multi-omics flow diagram demonstrating process of discovering and orthogonally validating gene marker panel. At each step, the number of marker genes is shown. **B)** Representation of the specimens for each step of the multi-omics validation. The GAstric Precancerous conditions Study (**GAPS**) is a prospective study, incorporating normal controls, low-risk precancerous lesions, high-risk gastric intestinal metaplasia (**GIM**), and early gastric cancer (**GC**). High-risk GIM lesions defined as Operative Link on GIM scores III or IV. GAPS specimens drawn equally from antrum (N=153) and body (N=150). A spatial transcriptomics cohort was derived from in-house sources. Bulk RNA-seq data from The Cancer Genome Atlas (**TCGA**) were used for validation. The scRNA-seq data was obtained both prospectively as well as through secondary analyses of published GIM and GC data sets. NAG, nonatrophic gastritis; CAG, chronic atrophic gastritis. *Immediately adjacent sections from the same tissue samples were used for spatial transcriptomics and smFISH.

The Gastric Precancerous Conditions Study (**GAPS**) is a prospective cohort of individuals undergoing endoscopy who are at increased risk for GC due to presence of symptoms (*e.g.,* dyspepsia, anemia), personal history (*e.g.,* GIM), or family history of GC. Enrolled subjects undergo biopsies according to the updated Sydney System, with standardized histologic assessment allowing for calculation of OLGIM stage^26^ and determination of Hp colonization. Sample-level phenotypic data and RNA sequencing metrics can be found in **Supplementary Table 1**.

The Cancer Genome Atlas Stomach Adenocarcinoma (**TCGA-STAD**) genomic dataset is comprised of GC samples which had not been previously treated by chemotherapy or radiation prior to genomic analysis.^27^ From these samples, we analyzed gene expression data from 180 intestinal-type GC primary tumors and 18 patient-matched tumor-adjacent controls. We obtained the de-identified patient clinical phenotype and RNA-seq results from Genomic Data Commons through TCGABiolinks^28^ R package. Tumor-level phenotypic data (*e.g.,* tumor location) is available in **Supplementary Table 2.**

We analyzed a scRNA-seq dataset for gastric pathology across Correa’s cascade (normal, NAG, CAG, GIM, and GC). This sample set constituted both *de novo* scRNA-seq data from prospectively collected samples along with public data sets. In total, the integrated scRNA-seq dataset comprised 40 biopsy samples from 26 patients: two normal controls, three NAG, three CAG, thirteen GIM, nine tumor-adjacent controls, and ten primary gastric tumors. Clinical phenotypic information (specimen location and histology), cell counts, and sequencing metrics are available in **Supplementary Table 3**.

For the spatial mapping and localization, we used formalin-fixed paraffin-embedded (**FFPE**) tissue specimens from five patients (three OLGIM II, one OLGIM III and one GC). The hematoxylin and eosin (**H&E**)-stained sections were manually annotated by an expert pathologist at the glandular level for regions of normal base, normal pit, metaplasia, dysplasia, and carcinoma. For spatial transcriptomics, unstained sections were placed on the Visium assay slide (10X Genomics) and stained with H&E followed by probe-based sequencing. For the single-molecule multiplex fluorescence *in situ* hybridization (**smFISH**) assays, we utilized unstained sections immediately adjacent (within 10 μm) to the Visium sections. Phenotypic description of the specimens used for spatial validation are available in **Supplementary Table 4**.

## RESULTS

### Overview of the multi-omics approach

An overview of the multi-omics pipeline is provided in **Figure 1A**, and a high-level description of the samples used for each step of the pipeline is provided in **Figure 1B**. In brief, this analysis included the following: i) discovery of a high-risk gene expression signature using RNA-seq data (N = 88 samples); ii) validation of the high-risk genes in a held-out cohort using RNA-seq data (N = 215 samples); iii) mapping of the high-risk genes to metaplastic foci using a spatial transcriptomics assay (N = 5 samples); iv) determining of the overlap of the high-risk GIM spatial signature with differentially expressed genes in intestinal-type GC samples with RNA-seq data (N = 198 samples); v) assigning the high-risk, spatially mapped genes to specific cell subpopulations using single cell RNA-seq (scRNA-seq) (N = 40 samples); and vi) validation of a subset of these genes at single cell resolution using smFISH (N = 5 samples).

### Gene expression analysis of high-versus low-risk GIM

For the GAPS-based marker discovery phase, a detailed summary of the cohort’s demographic, clinical and histologic characteristics are provided in **Table 1**. The cohort was highly diverse with multi-ethnic representation. Samples originated from a high proportion of Hispanic (23.9%), Asian (42.9%), and foreign-born (62.6%) subjects. When assessing OLGIM stages, 56.4% were OLGIM stage 0 (no GIM), 16.6% OLGIM I, 13.5% OLGIM II, 9.2% OLGIM III, and 4.2% OLGIM IV.

**Table 1:** Clinical and Histopathologic Attributes of Hp-negative Cohort.

|  |  |  |  |
| --- | --- | --- | --- |
| Enrolled Subjects (N=163) |  | Unique Gastric Biopsies (N=303) |  |
| Characteristic | Frequency (%) | Finding | Frequency (%) |
| Age |  | All biopsies |  |
| <50 | 40 (24.5) | Normal/NAG | 204 (67.3) |
| 50-69 | 90 (55.2) | CAG | 99 (32.7) |
| >70 | 33 (20.2) | GIM severity | 96 (31.7) |
| Female | 105 (63.1) | Mild | 42 (13.9) |
| Race/Ethnicity |  | Moderate | 31 (10.2) |
| Non-Hispanic White | 37 (22.7) | Severe | 23 (7.6) |
| Black | 1 (0.6) | Antrum | (N=153) |
| Hispanic | 39 (23.9) | Normal/NAG | 92 (60.7) |
| East Asian | 70 (42.9) | CAG | 61 (39.9) |
| Other | 16 (9.8) | GIM severity | 60 (39.2) |
| Foreign born | 102 (62.6) | Mild | 26 (17.0) |
| Family history* | 16 (9.8) | Moderate | 20 (13.1) |
| Proton pump inhibitor use | 53 (20.4) | Severe | 14 (9.2) |
| OLGIM stage** |  | Body | (N=150) |
| 0 (no GIM) | 92 (56.4) | Normal/NAG | 112 (74.7) |
| I (lowest) | 27 (16.6) | CAG | 38 (25.3) |
| II | 22 (13.5) | GIM severity | 36 (24.0) |
| III | 15 (9.2) | Mild | 16 (10.7) |
| IV (highest) | 7 (4.2) | Moderate | 11 (7.3) |
|  |  | Severe | 9 (6.0) |

We used conventional RNA-seq to analyze 303 gastric specimens (153 antrum, 150 body) originating from 163 unique individuals from GAPS (specimen-level data on histopathology and sequencing metrics are provided in **Supplementary Table 1**). The cohort was separated into a discovery set of 88 GIM samples (22 high-risk, 66 low-risk) from 46 patients and a held-out validation set of 215 GIM samples (22 high-risk, 193 low-risk) from 115 patients. Prior to differential expression analysis, we performed unsupervised clustering through i) hierarchical clustering with Pearson correlation distance, as well as ii) principal components analysis (**Supplementary Figure 1**) to confirm preferential grouping of high-risk and low-risk samples. Subsequently, we conducted differential expression analysis with limma-voom,^29^ utilizing a factorial design strategy (**Supplementary Figure 2**).

### Discovery of genes associated with high-risk GIM

From the discovery data set of 88 GIM samples, we identified a preliminary set of 399 genes that were differentially expressed in the high-risk patients in both the body and antrum of the stomach (**Supplementary Figures 3A – C; Supplementary Table 5**). Expression differences were based on a fold-change cutoff ≥ 1.25 in the linear scale at a false discovery rate-adjusted p-value < 0.05. Likewise, we excluded any genes which differential expression profile differed significantly (significant interaction term) between antrum and body (**Supplementary Figures 3D**). These genes included established markers of intestinal metaplasia (*i.e.*, *CDX1, CDX2, OLFM4, ACE2*) and gastric epithelial cells (*i.e.*, *PGC, CCKBR*).

Next, we conducted weighted gene co-expression network analysis (**WGCNA**)^30^ to determine groups of co-expressed genes, otherwise referred to as gene modules. Gene modules were summarized using the module eigengene (the first principal component of the gene expression levels within a module). We leveraged this information to determine whether the expression levels of the genes from each module were directly or inversely correlated with high-risk GIM (*i.e.*, module-trait relationships) in an independent approach, separate from the differential expression analysis. We selected two modules associated with high- and low-risk samples (**Supplementary Figure 4A**). Using hierarchical clustering with Pearson correlation distance, we demonstrated that genes form these modules are informative of high-risk gastric cancer precursors (**Supplementary Figure 4B**), whereas five other modules were not informative (**Supplementary Figure 4C**). We intersected genes from the two informative WGCNA modules and the differential expression analysis, resulting in a refined set of 314 genes that were: i) differentially expressed in high-risk samples from both anatomic regions, and ii) co-expressed in gene modules associated with high-risk stages (**Figure 2A, Supplementary Figure 4D, Supplementary Table 5**). Using these 314 genes, most of the high-risk samples clustered distinctly and separately from the low-risk group, regardless of the anatomic origin of the biopsy (antrum or body). By contrast, among the low-risk samples (OLGIM 0, I or II), gene expression showed clustering for either the antrum or body (*top dendrogram*). This result indicates that the antrum and body are transcriptionally distinct entities. However, once advanced GIM develops, a specific expression profile emerges regardless of anatomic origin.

**Figure 2:**
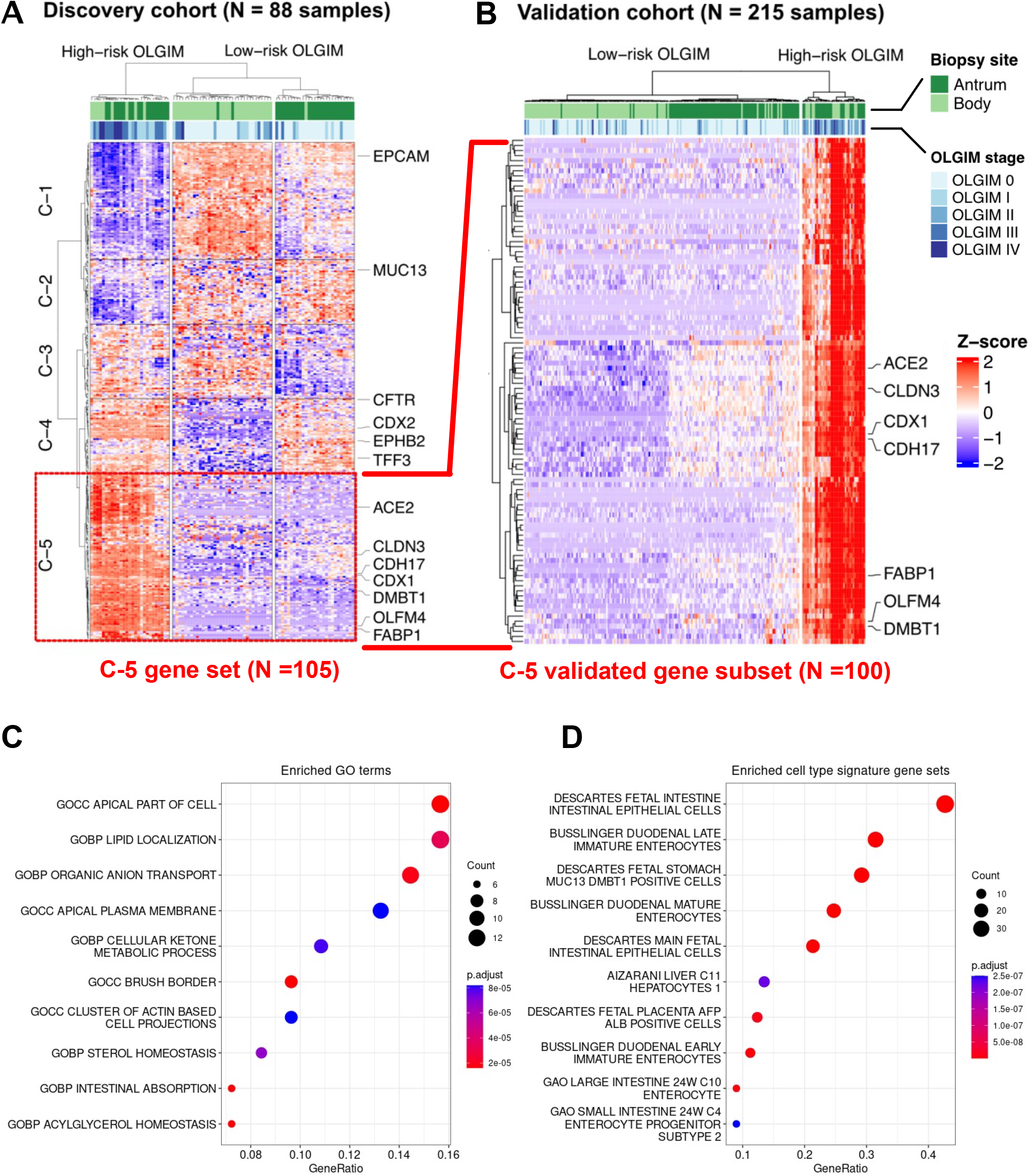
Discovery and validation of the high-risk expression signature. **A)** Heatmap and hierarchical clustering of differentially expressed and co-expressed genes from the discovery cohort of 88 samples, 22 high-risk (defined as OLGIM III-IV) and 66 low-risk (defined as OLGIM 0-II). Most of the high-risk samples clustered distinctly and separately from the low-risk group, regardless of the anatomic site of the biopsy (top dendrogram). A set of genes were found to be both differentially expressed between high- and low-risk samples. Cluster-5 (C-5) represents 105 genes which were selectively upregulated in their expression only in high-risk samples, regardless of anatomic location. **B)** We found 100 genes from C-5 to be differentially upregulated in the validation cohort (22 high-risk and 193 low-risk samples), confirming a robust signature for high-risk GIM which is agnostic of location. Dotplot depicting over-representation analysis results of these 100 genes: **C)** gene ontology terms are enriched with intestinal processes (*e.g.,* brush border, intestinal absorption); **D)** cell type signature gene sets are enriched for mature and immature/fetal intestinal cell types.

From this subset of 314 genes, we identified five discrete expression clusters that were labeled C-1 through C-5 (**Figure 2A**, *side dendrogram*). Cluster C-5 represented a subset of 105 genes which were overexpressed in high-compared to low-risk samples, with the highest Z-scores. Importantly, the C-5 cluster was independent of the anatomic location. This gene set included established GIM markers such as *CDX1*, *FABP1* and *ACE2* (**Supplementary Table 5**). In addition to markers of mature enterocytes (*ANPEP*, *CDH17*),^31,32^ we also found markers of intestinal stem cells (such as *OLFM4*),^33^ and markers of transit amplifying cells (*DMBT1*) in this cluster.^34^

### Validating the genes associated with high-risk GIM

Next, we validated the results from the discovery analysis, performing differential expression analysis on the held-out, independent validation set of 215 GIM samples. We focused on the C-5 cluster, given that overexpressed genes provide an opportunity for additional experimental validation. For the 105 genes identified from the C-5 cluster in the discovery set, we found a striking 100 out of 105 genes (95.2%) consistently overexpressed (linear fold-change ≥ 1.25 and FDR-adjusted p < 0.05) among the validation set’s high-risk samples (**Figure 2B**). The full gene list is provided in **Supplementary Table 6**. To characterize the functional pathways and cellular associations of these 100 genes, we conducted over-representation analyses with clusterProfiler.^35,36^ We selected gene sets relative to gene ontology and cell types from the MSigDB database.^37–39^ Enriched gene ontology terms (**Figure 2C**) included intestinal absorption (*SLC2A5*, *ABCG8*, *ABCG5*, *MOGAT2*, *PRAP1*, *FABP1*) and presence of a brush border (*ACE2*, *SLC28A1*, *SLC2A2*, *MME*, *SLC6A19*, *SLC7A9*, *MTTP*, *MYO7B*) among other intestinal-related processes. Consistent with these findings, we found enrichment of certain cell lineage gene sets (**Figure 2D**) including mature (*SLC2A5*, *APOC3*, *ACE2*) and fetal enterocytes (*LRRC19*, *CELP*, *RBP2*), as well as immature enterocytes (*DMBT1*, *CPS1*). A comprehensive listing of enriched gene ontology terms and cell lineage gene sets are available in **Supplementary Tables 9 and 10.**

### Spatial transcriptomics maps the high-risk expression signature to metaplastic gastric foci

Next, we used a spatial expression assay (Visium, 10X Genomics) to map the genes of the high-risk expression signature to GIM regions. Spatial transcriptomics were applied to five GIM samples with extensive histopathology annotation and all of them included regions of i) normal base, ii) normal pit or iii) metaplastic foci. An example of one GIM sample (**P09788**) is shown in **Figure 3**. The aggregated spots per area for each sample were used to conduct a differential expression analysis comparing separate regions of different histologic annotation, similar to an *in-silico* tissue microdissection. Among all five GIM, we determined that 458 genes were significantly upregulated in regions of metaplastic foci compared to both normal gland base and pit (‘pseudo-bulk’ analysis, **Supplementary Table 7**).

**Figure 3:**
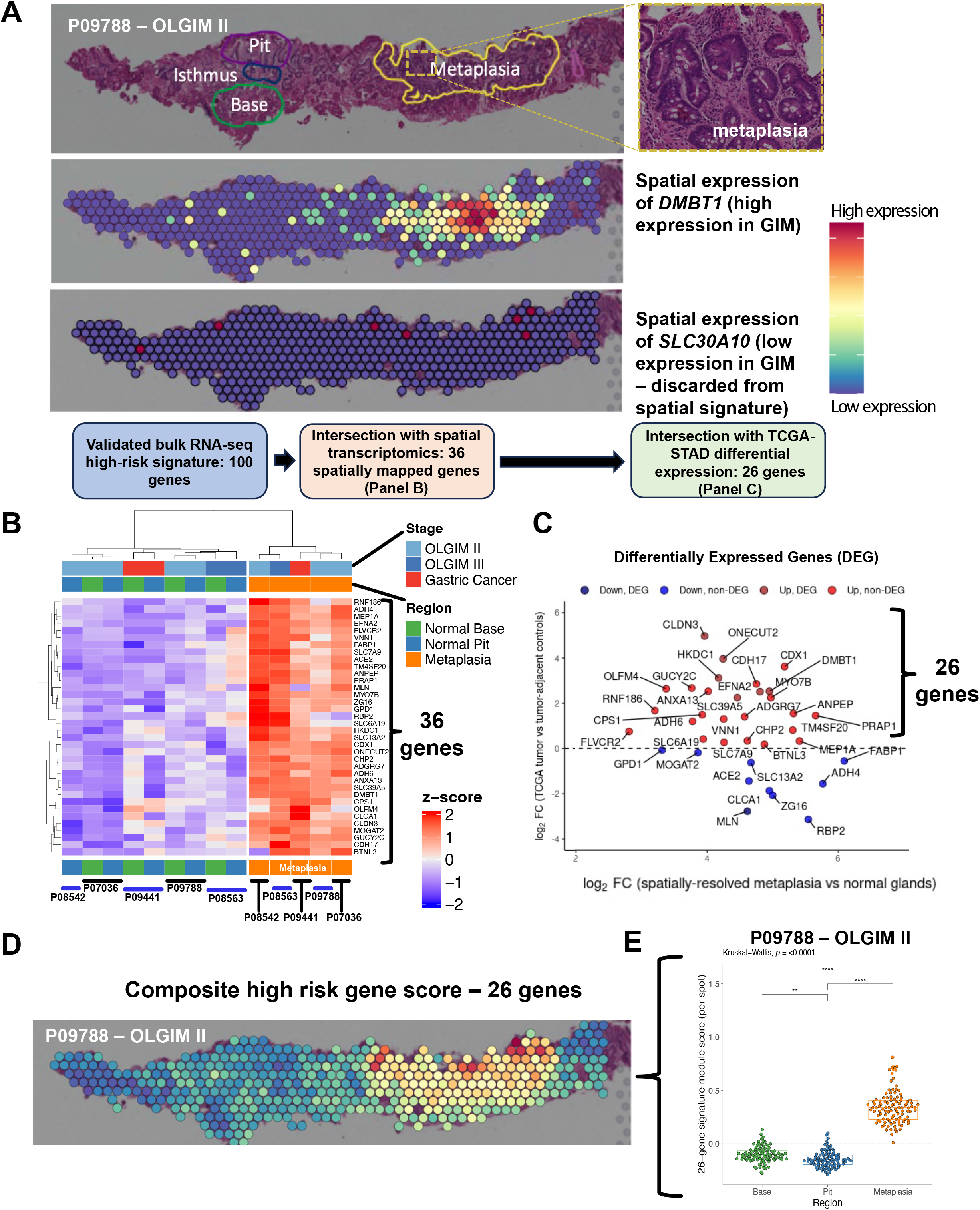
Spatial resolution of the high-risk signature. **A)** An example of the expression profile of *DMBT1* upon a Visium slide annotated by a pathologist for areas of normal glandular architecture (base and pit) and metaplasia. *DMBT1* is shown as an example of a spatially resolved gene mapping to pathologist-annotated metaplasia, whereas *SLC30A10* is shown as an example of a gene not mapping to metaplasia and, thus, discarded form the spatially-resolved signature. **B)** Heatmap depicting 36 differentially expressed genes from spatial pseudobulk analysis that overlapped with bulk RNA-seq signature. **C)** Scatter plot showing log_2_ fold-change of 36 genes from spatial pseudobulk analysis (X-axis) and log_2_ fold-change from TCGA-STAD RNA-seq analysis. Twenty-six genes overexpressed in both analyses are shown in red. **D)** Spatial mapping of the refined 26-gene signature onto Visium spots. **E)** Comparison of 26-gene signature between metaplastic foci vs normal stomach base or pit (Kruskal-Wallis and Dunn test FDR-adjusted *p* < 0.001). Note: each Visium spot is 55 µm in diameter, with 100 µm distance between the center of adjacent spots.

Next, we intersected the 100 genes from the validated expression signature as previously described with the 458 genes that were mapped using the spatial transcriptomic assay. Notably, from the validated expression signature, 36 out of 100 (36%) genes were expressed specifically in regions of the metaplastic glands (*e.g.*, *DMBT1*, **Figure 3A**; other spatially resolved genes in **Figure 3B**). Genes from the signature that did not map to metaplasia were discarded (*e.g.*, *SLC30A10*, **Figure 3A**). Overall, this result identified a subset of 36 high-risk differentially expressed genes that mapped to pathologist-annotated regions of metaplasia.

### The high-risk expression, spatially mapped signature’s association with gastric cancer

We determined how many of the 36 spatially mapped high-risk genes were also differentially expressed in the intestinal subtype of GC. This step of the analysis used RNA-seq data from the TCGA-STAD cohort. We conducted differential gene expression analysis between 180 gastric cancers, all being of the intestinal subtype, and 18 matched tumor-adjacent gastric tissues. We compared the fold-change from the TCGA analysis vs the fold-change from the spatial gene expression analysis for the 36 genes (**Supplementary Table 8**). Twenty-six genes overlapped between those which were significantly overexpressed in high-risk GIM (relative to low-risk GIM), localized to metaplastic foci, and were consistently upregulated in GC (**Figure 3C**).

We quantified the expression of these 26 genes in gastric metaplastic foci using a composite signature score using Seurat’s AddModuleScore function.^40,41^ This algorithm calculates the average expression levels of a gene set cluster when compared to the aggregated expression of a control gene set. We used this method to compare expression among regions of normal gland base, pit, and metaplasia. The 26-gene score was significantly higher among the Visium spots mapping to metaplastic foci for each of the five spatial samples compared to areas with normal stomach base or pit (Kruskal-Wallis and Dunn test FDR-adjusted *p* < 0.001) (**Figure 3D** **and Supplementary Figure 5**). Overall, this set was highly specific for metaplasia and did not map to any other normal gastric regions, as well as tumor regions (moderate signal) in one GC sample. The gene signature included established markers for immature intestinal lineages (*OLFM4*, *DMBT1*)^33,34^ and markers for mature enterocytes (*ANPEP*, *CDH17*)^31,32^ (**Supplementary Table 8**).

### The 26 gene high-risk spatial signature is expressed in goblet cells, enterocytes, and intestinal stem-like cells

For the next step, we used scRNA-seq results to determine which cell types expressed the 26 gene high-risk signature. This analysis used a set of 40 specimens of both GC and precancerous lesions. The results provided the assignment of the high-risk expression signature to specific single cells and lineages. The joint data set contained a total of 116,643 single cells. From this data set we identified seven major cell lineages: epithelial, T cells, B cells, stromal cells, endothelial, myeloid and mast cells. Using the Seurat AddModuleScore function, we determined that the 26 gene signature across all cells was significantly higher among epithelial cells (**Supplementary Figure 6**).

We characterized the expression features of the epithelial subset (**Figure 4A**). Cell identity of this subset was determined using reference mapping on a cell atlas encompassing normal stomach and duodenum. Cell types included i) normal gastric lineages (chief, parietal, endocrine cells), ii) mature intestinal cells (goblet cells, enterocytes), and iii) a broad class of immature intestinal cells (duodenum stem cells, duodenum differentiating stem cells, and duodenum transit amplifying cells). We call these immature cells collectively “intestinal stem-like cells”. We found the 26-gene signature to be significantly enriched in enterocytes and intestinal stem-like cells (**Figure 4B** **and 4C**). Conversely, this 26-gene signature was nearly absent among all normal gastric lineages (Dunn test FDR-adjusted *P* ≤ 0.001, **Supplementary Table 11**).

**Figure 4:**
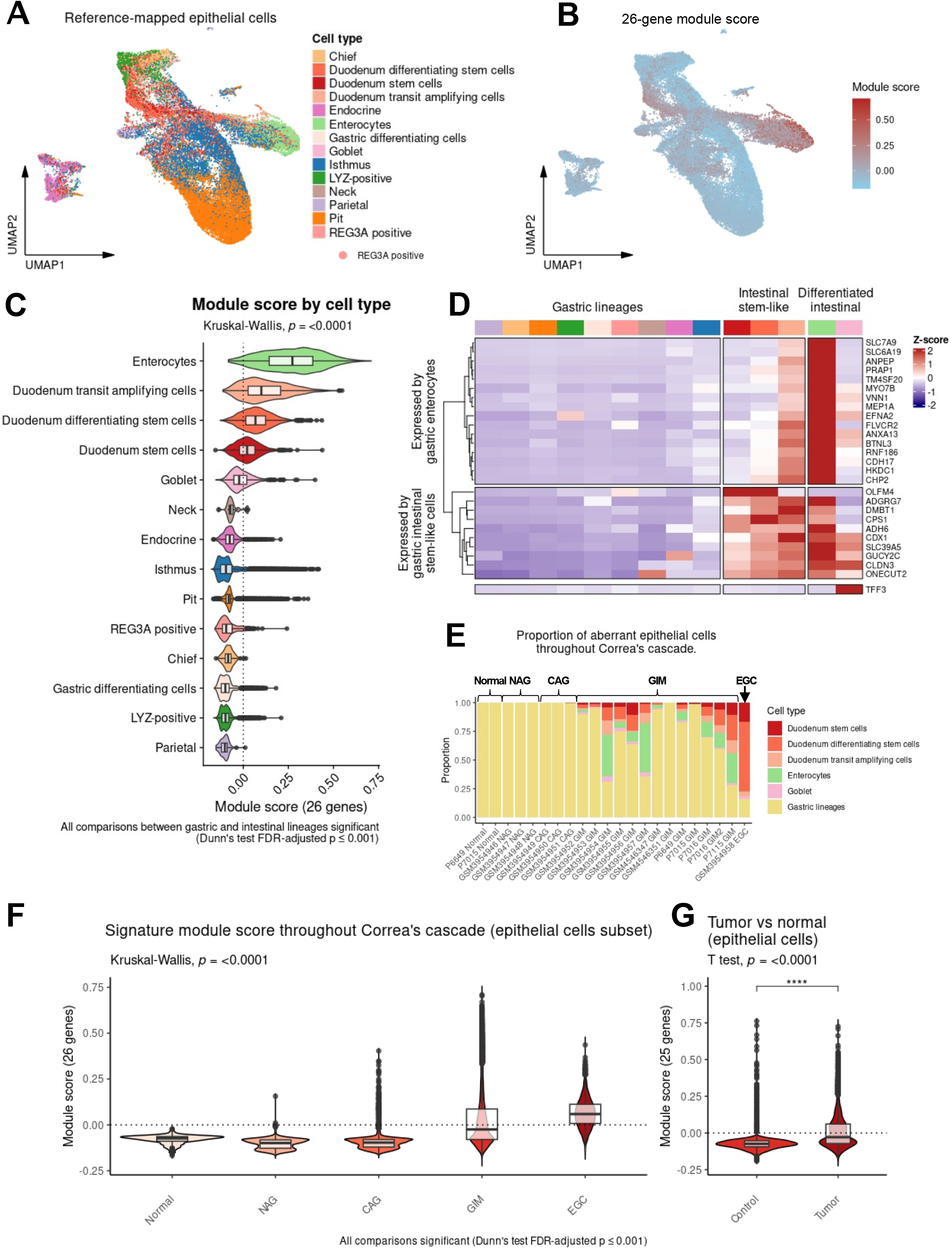
Single-cell identification of cell types expressing the high-risk signature. **A)** UMAP plot showing reference-mapped epithelial cells. **B)** UMAP plot showing module score. **C)** Comparison of the module score between cell types. **D)** Heatmap showing the scaled expression of the 26 genes by cell type. The gene dendrogram shows genes expressed primarily by “gastric” enterocytes and another expressed predominantly by “gastric” intestinal stem-like cells. Cells identified as goblet cells expressed TFF3. **E)** Stacked bar plots depicting the proportion of cell types per sample, ordered by stage of Correa’s cascade. Gastric lineages are aggregated into a single class. **F)** Comparison of module score across Correa’s cascade (p<0.001 for all comparisons). **G)** Comparison of the module score between GC and tumor-adjacent control tissues (p<0.0001).

Next, we examined the expression of each gene scaled across the epithelial cell types to highlight which cells displayed the highest levels of expression. As a specific marker for goblet cells, we also included the gene *TFF3*.^42,43^ We performed hierarchical clustering of the 26 genes and *TFF3* from the signature. The gene dendrogram revealed two distinct groups of genes, one that is expressed primarily by differentiated “gastric” enterocytes, and one expressed by “gastric” intestinal stem-like cells, given their gastric localization (**Figure 4D**). The duodenum stem cells were characterized by high expression of *OLFM4*, a known intestinal stem cell marker,^32^ as well as *ADGRG7, CPS1, ADH6, SLC39A5, GUCY2C, CLDN3, AND ONECUT2.* Duodenum differentiating stem cells mostly expressed *OLFM4* and *CPS1* but also expressed *ADGRG7, DMBT1, ADH6, CDX1, SLC39A5, GUCY2C, CLDN3, AND ONECUT2*. Duodenum transit-amplifying cells, an undifferentiated population transitioning between stem cells and differentiated intestinal cells, showed reduced expression of *OLFM4*, reduced expression *CPS1,* and expression of *DMBT1*, an established marker that has been noted in other reports.^34,44^

We observed that many of the genes comprising the intestinal stem-like cell markers group were carried over to differentiated enterocytes. Further, some of these genes (*ADGRG7, ADH6, SLC39A5, GUCY2C* and *CLDN3*) showed increased expression among enterocytes, suggesting that these genes are expressed at progressively increasing levels throughout the differentiation process. We determined that *TFF3* was predominantly expressed among goblet cells.

Next, we analyzed the proportion of different cell lineages throughout Correa’s cascade (**Figure 4E**). Importantly, normal, NAG, and CAG gastric tissue samples did not have mature or immature intestinal cell lineages. In contrast, GIM was characterized by both the presence of mature (goblet cells and enterocytes) and immature intestinal cells. Moreover, GC was characterized by a significant enrichment of immature intestinal lineages and substantial loss of differentiated goblet cells and enterocytes. These results suggest that GIM is characterized by the onset and expansion of both mature and immature intestinal cell types. The continued expansion of this immature population may be an important indicator of progression to intestinal GC. Our results support this interpretation of the results. Namely, we observed that the 26-gene signature score increased with the progressive stages of Correa’s cascade (Dunn’s test FDR-adjusted *P* ≤ 0.001 for all comparisons; **Supplementary Table 12** and **Figure 4F**). In a separate analysis, we analyzed the signature score between GCs and patient-matched tumor adjacent control tissues from a previous publication^45^ (**Figure 4G**). As expected, we found the 26-gene signature to be highly-enriched in tumor cells relative to adjacent non-tumor, normal gastric tissue (Welch’s T test *P* < 0.0001).

### Single-molecule fluorescent *in situ* hybridization reveals intestinal stem-like cells in the isthmic/crypt region of metaplastic glands

We conducted a higher resolution spatial analysis to identify single cells expressing components of the high-risk signature. The smFISH assay uses *in situ* RNA hybridization to visualize the spatial location of specific expressed genes at single-cell resolution and enables comparisons to standard histopathology. This technique allows one to assess multiple genes’ spatial relationship and expression among the individual cells from a given tissue section. Based on the scRNA-seq results that defined the aberrant intestinal stem cell populations, we selected eleven of the signature genes for smFISH: *ANXA13*, *HKDC1*, *OLFM4*, *DMBT1*, *CPS1*, *SLC39A5*, *ANPEP*, *ONECUT2*, *CDH17*, *CLDN3* and *CDX1*. These genes included markers for intestinal stem-like cells (*OLFM4, CPS1, DMBT1)*, enterocytes (*HKDC1, ANPEP, CDH17, CLDN3, ANXA13*), or were expressed across all intestinal lineages (*CDX1, SLC39A5, ONECUT2*). We also included *TFF3* as a specific goblet cell marker. After imaging, the results were compared to the matching H&E images with pathology interpretation.

We performed smFISH (RNAscope HiPlex) on tissue sections adjacent to those analyzed in the prior spatial transcriptomics assay. We identified several distinct cellular compartments which occurred within metaplastic tissue (**Figure 5, Supplementary Figures 7-11**). The first compartment consisted of mature or differentiated intestinal cells and were characterized by strong expression of *TFF3* (goblet cells) and moderate signal of *ANPEP* (enterocytes). A second compartment consisted of cells which strongly expressed stem-like markers (O*LFM4, DMBT1* and *CPS1*); these columnar cells were characterized by high nuclear-to-cytoplasm ratio, were located near crypt regions of metaplastic glands, and were mutually exclusive in space to the mature markers. These results provide additional evidence confirming the presence of intestinal stem-like cells as previously identified in the scRNA-seq results. There were some genes (*ONECUT2* and *HDKC1*) which were ubiquitously expressed in both mature and stem-like cells, although expression of these genes was higher among stem-like cells. Notably, none of the selected genes were expressed in the normal gastric glands across any of the samples. The results from smFISH provided single cell spatial resolution and confirmed the presence of distinct cellular compartments within metaplastic glands consisting of either mature or immature intestinal lineages. Also, this analysis revealed the spatial relationship between these two distinct cellular populations relative to glandular architecture.

**Figure 5:**
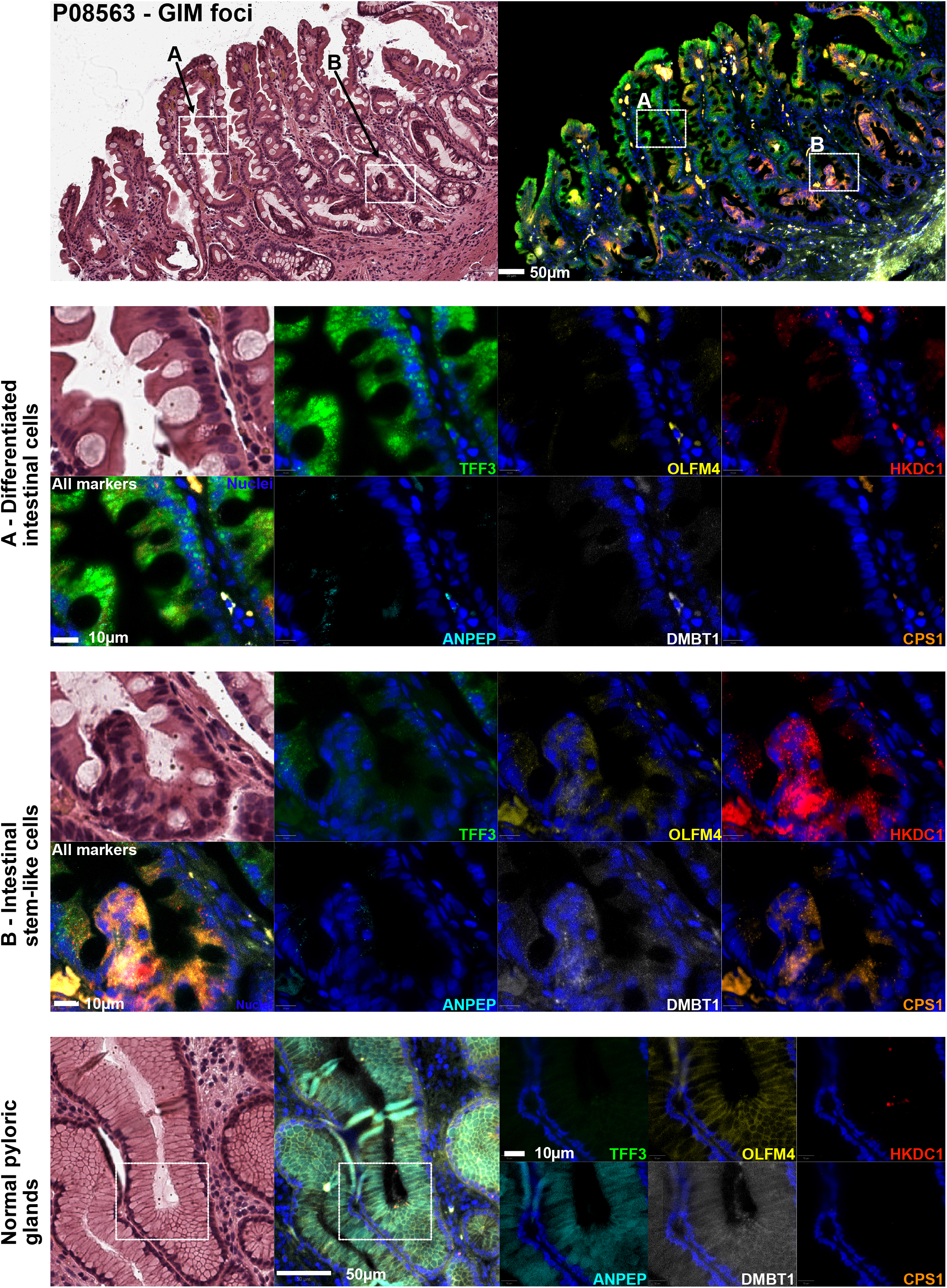
Single-molecule fluorescence in situ hybridization (smFISH) of gastric intestinal metaplasia. Representative region showing H&E staining and smFISH for 6 genes in a GIM foci: *TFF3, HKDC1, DMBT1, OLFM4, CPS1,* and *ANPEP*. **A)** Areas of mature intestinal cells demonstrated robust expression of *TFF3* (goblet cells) and *ANPEP* (enterocytes). **B)** Other genes (*OLFM4, DMBT1, CPS1, HKDC1*) localized to columnar cells near the isthmic/crypt regions in the metaplastic glands (intestinal stem-like cells). The expression of these genes was mutually exclusive, in space, from the mature markers. **C)** Representative normal gland showing that genes from the signature are not expressed by normal gastric lineages.

In the tumor sample, we also identified distinct compartments of well-differentiated cells that expressed *TFF3.* Notably, poorly differentiated regions of the tumor showed high expression of *HKDC1* and *OLFM4.* In addition, defined cell clusters within the poorly differentiated regions of the tumor co-expressed *CDX1, OLFM4, CPS1, DMBT1* and *HKDC1* (**Supplementary Figure 11**).

## DISCUSSION

In this study, we analyzed a Hp-negative cohort of individuals with pathology across the gastric precancerous spectrum. Using integrated bulk gene expression, spatial transcriptomics, scRNA-seq, and smFISH, we identified a highly refined signature of 26 genes which characterized high-risk gastric precursors. Notably, this set of genes was i) associated with advanced OLGIM stages, ii) localized to metaplastic glands on histopathology, iii) expressed by aberrant epithelial cells not typically found in healthy gastric tissue, iv) differentially expressed from metaplasia to intestinal-type GC, and v) distinguished between mature cells (goblet cells and enterocytes) and stem-like cells in metaplastic foci.

While this high-risk gene expression signature is characterized by both markers of mature enterocytes and stem-like cells, we found that the more advanced lesions had greater expression of immature intestinal markers. This progression was characterized by increased expression of intestinal stem cell markers such as *OLFM4*,^33^ as well as markers of transit amplifying cells (*DMBT1*)^34^. These markers were absent from both normal differentiated gastric tissues, as well as gastric stem cells. Collectively, these results point to intestinal stem-like cells playing an important, constitutive role in the biology of advanced preneoplasia. A recent longitudinal study from a Singaporean consortium also found changes in the intestinal stem-like cell compartment to be associated with risk for GC.^46^ In addition, another recent report demonstrated phenotypic mosaicism in GIM, by which individual GIM cells co-express both intestinal and gastric markers.^47^ This interesting finding strongly suggests that the intestinal cells identified by our studies also possess gastric transcriptional properties, substantiating the terminology of “gastric enterocytes” or “gastric intestinal stem-like cells”. Future efforts should be made to fully characterize these cell populations. Our data lend further credence to the hypothesis that terminally-differentiated epithelial cells (such as enterocytes and goblet cells) may simply be passive bystanders harboring genetic alterations already present in genetically unstable stem-like cells, the latter of which have the potential to become the true carcinogenic precursors.^48^

Among the genes from the high-risk signature, some correspond to known markers for intestinal stem-like cells, including *OLFM4* and *DMBT1*. Other genes in the signature are established markers for mature enterocytes. For example, *ANPEP* encodes aminopeptidase N, an enzyme found in the apical membrane of mature enterocytes. An early report suggested that leucine aminopeptidase activity was highly specific to metaplastic zones within the human stomach examined microscopically.^31^ *CDH17* is a membrane-associated enterocyte marker that has been found expressed in >60% of GCs, with greater expression specifically in intestinal-type GCs.^32^

In our study, we found that *CPS1* to localize selectively to intestinal stem-like cells. Interestingly, *CPS1* is an enzyme of the urea cycle and has previously been associated with GIM.^49^ Thus far, *CPS1* has not been associated with stem cell biology. However, a recent report in lung cancer suggests that *CPS1* may be crucial for pyrimidine maintenance and DNA synthesis in *KRAS* mutant cells.^50^ Notably, *KRAS* was one of the driver oncogenes previously identified,^46^ suggesting that there may be a relevant and novel role for *CPS1* as a source supply of pyrimidines in the context of DNA synthesis among replicating precancerous cells.

*HKDC1* is a hexokinase and has not been previously described in gastric preneoplasia. Two independent *in vitro* studies have evaluated the role of *HKDC1* in gastric cells. The first recently found that *HKDC1* can promote chemoresistance, proliferation and invasion, and epithelial-to-mesenchymal transition through induction of NF-κB.^51^ The second study suggests *HKDC1* may be pivotal for glycolysis and proliferation in AGS and MKN-45 GC cells.^52^ Supporting a relevant role for *HKDC1* in carcinogenesis, this gene has been found to promote lung,^53^ breast,^54^ and biliary^55^ cancers. Moreover, deletion of *HKDC1* inhibits proliferation and tumorigenesis in mice.^56^ Our findings suggest *HKDC1* may be a novel marker for advanced gastric preneoplasia.

Spatial co-expression of *HKDC1* and *CPS1* with *CDX1, OLFM4* and *DMBT1* in GIM samples, as well as GC, provide strong evidence of the potential role of these cells in metabolic reprogramming of GC precursor lesions. This is highly consistent with the single-cell expression profiles. Further studies are warranted to establish their role in gastric preneoplasia.

Most prior molecular and genomic studies of the GIM microenvironment have focused on populations with moderate-to-high Hp prevalence and high GC incidence.^22,46,57^ Our study addresses a gap in the literature by providing needed mechanistic data on advanced GIM in a relatively low-risk population common to many regions of North America and Europe. An additional motivation for developing an Hp-negative biomarker is that almost all patients diagnosed with GIM have already undergone Hp eradication therapy — that is to say, by the time such patients come to clinical attention they are functionally Hp negative. The ongoing carcinogenic potential of Hp-negative GIM may in part be explained to the establishment of clonal stem-like cell lineages, as suggested by this and other studies.^46,57^

Our findings have public health significance. As only a small fraction (<5%) of patients with GIM will progress to GC over long-term follow-up,^9–13^ indefinite endoscopic surveillance of these patients may lead to unnecessary cost, medical risk, and anxiety. It has been suggested that combined clinical-genomic models may outperform clinical-only models in predicting individuals most at risk for GC progression.^22,46^ This new high-risk signature may thus serve an important risk-stratification purpose in individuals diagnosed with GIM.

Our study benefited from the prospectively collected specimens from the GAPS study which have the most extensive clinical annotation of all the data sets. As a result, we had histopathologic scoring by OLGIM. In contrast, public data sets of GIM were available only in broader classes of Correa’s cascade (*e.g.*, NAG vs CAG vs GIM). Additional steps with the cohorts that did not have as extensive clinical annotation (i.e., GIM from public scRNA-seq results) were used for marker refinement, and to demonstrate that a consistent trend toward enhanced expression was evidenced throughout Correa’s cascade.

Spatial validation assays (both Visium and smFISH) are newly-emerging approaches. These results provided the high resolution cellular imaging of the GIM spatial microenvironment. Moreover, each sample in the validation steps required detailed annotation by an expert pathologist; as such, each slide represents hundreds of independent, phenotyped data points on which analysis was performed.

Our study was cross-sectional in nature and did not contain longitudinal data on GIM progression. Our future studies will be focused on establishing causal inference through prospective longitudinal cohorts. From these type of samples, we will conduct future studies in which we predict histologic progression toward neoplasia in a cohort of patients with preneoplastic lesions.

## CONCLUSION

Leveraging multiple independent cohorts, we utilized integrated transcriptomic approaches incorporating both spatial and single-cell methods to further characterize the molecular and cellular origins of high-risk GIM. We identified a discrete set of 26 genes which are associated with higher OLGIM stages, localize spatially to metaplastic foci, are expressed by aberrant epithelial cells, are differentially expressed in intestinal-type GC, and characterize both mature and immature intestinal cells. We find that with increasing histologic severity, the expression of intestinal stem-like cell markers increases. These data hold important future implications for future cancer interception.

## Acknowledgements

We thank all study participants for their contribution to this study. We also thank Advanced Cell Diagnostics for the RNAScope HiPlex reagents, received as part of the spatial grant program to IAW.

## Authors’ contribution

RJH co-led manuscript writing, co-led conception and design, and co-led specimen collection. IAW co-led manuscript writing, led bioinformatics and statistical analysis, drafted figures and co-led interpretation of data. AS contributed to statistical analysis, interpretation of data, and figure creation. AS performed experimentation including scRNA-seq and data analysis. MVS performed specimen collection, specimen processing, and table generation. SG performed bioinformatics analysis. RM performed experiments in support of smFISH. AA performed data collection, specimen collection, and specimen processing. XB performed statistical analysis. JS interpreted and annotated biopsies. QN assisted in the bioinformatics analysis. MRA contributed to conception, design, and image generation. JHH co-led conception and design, co-led specimen collection. HPJ co-led conception and design and provided overall administrative oversight over the project. All authors participated in drafting the manuscript for important intellectual content and gave approval for publication.

## Ethics approval

The collection of clinical data and biological specimens from GAPS, the spatial cohort, and in-house scRNA-seq samples were approved by the Stanford University Institutional Review Board (45077).

## Data availability statement

Data from this study are available in phs001818.v1.p1. Downloaded datasets are available under accession numbers GSE134520, GSE150290, and PRJNA678538. The code used in this study can be accessed at https://github.com/sgtc-stanford/GIM-spatial.

