## Supplemental Methods and Figures for "A spatially mapped gene expression signature for intestinal stem-like cells identifies high-risk precursors of gastric cancer"

#### SUPPLEMENTARY METHODS

##### Overview of the data sets

The study used samples and clinical data from four sources: 1) a prospective study entitled the GAstric Precancerous conditions Study (**GAPS**, NCT04191551), that includes patients undergoing endoscopic resection at Stanford Health Care; 2) RNA-seq data from The Cancer Genome Atlas stomach adenocarcinoma cohort (**TCGA-STAD**);[1] 3) a scRNA-seq dataset comprised of both previously published and *de novo* data derived from patients with normal histology, non-atrophic gastritis (**NAG**), chronic atrophic gastritis (**CAG**), gastric intestinal metaplasia (**GIM**), and gastric cancer (**GC**); 4) as well as a spatial validation cohort composed of pathologist-annotated slides from Stanford Health Care patients.

**1) GAPS**: GAPS is a prospective cohort of individuals undergoing endoscopy who are at increased risk for GC due to presence of symptoms, personal history, or family history of GC. Participants were between the ages of 35 and 84 and were enrolled for the following indications: abdominal pain, dyspepsia, iron deficiency anemia, *Helicobacter pylori* (**Hp**) assessment, surveillance of known GIM, and evaluation for family history of GC. Biopsy samples from both the gastric antrum and gastric body were obtained during endoscopy following the updated Sydney System biopsy protocol and examined by a pathologist to assess precancerous lesions, operative link (**OLGIM**) staging,[2] and presence of Hp. Two additional mucosal biopsies were frozen and stored at -80 °C for sequencing and downstream analyses. Presence of Hp was determined by IHC staining using the SP48 monoclonal primary antibody (Ventana Medical Systems, Tucson, AZ). In addition, Hp-negativity was confirmed through detection of Hp 16s sequencing reads of the V3 and V4 regions in the training set. For pathology evaluation and OLGIM staging, the presence of metaplasia was categorized as none [0% involved], mild [1-30%], moderate [31-60%], or severe [>60%] for each of the biopsies from the Sydney protocol. The OLGIM score was then calculated based on a composite of scores from the antrum and body. Sample-level phenotypic data and mapped sequence reads can be found in **Supplementary Table 1** (**ST1**).

2) **TCGA-STAD:** The **TCGA-STAD** genomic dataset comprises 295 GC samples which had not been previously treated by chemotherapy or radiation prior to genomic analysis.[1] We selected a subset of 180 intestinal-type GC primary tumors and 18 patient-matched normal gastric tissue. The de-identified patient clinical phenotype and processed RNA-seq counts came from the Genomic Data Commons can be accessed through their online data portal (https://gdc.cancer.gov/).[3] We downloaded the data through TCGABiolinks[4] in R statistical programming environment. Tumor-level phenotypic data (*e.g.,* tumor location) can be found in **Supplementary Table 2.**

3) **Single-cell RNA-seq datasets:** We analyzed an integrated **scRNA-seq dataset** for gastric samples with pathology across Correa's cascade (normal, NAG, CAG, GIM, and GC). This sample set constituted both *de novo* scRNAseq data from prospectively-collected samples along with four public data sets. We generated *de novo* data from two patients for this study: one patient with matched biopsies from GIM and normal glands, and another patient with matched GIM biopsies from the antrum and body. For the GC samples, we utilized previously reported scRNA-seq from both the primary tumor along with patient-matched tumor-adjacent control tissues (18 biopsies from 7 patients).[5] We used scRNA-seq data from patient-matched GIM with normal gland biopsies from the same study;[5] one GIM;[6] twelve samples from a large public scRNA-seq data corresponding to three NAG, three CAG, six GIM and one early GC biopsy (GEO accession number GSE134520);[7] and two individuals with GIM (GEO accession number GSE150290).[8] In total, the integrated scRNAseq dataset comprised 40 biopsy samples from 26 patients: two normal controls, three NAG, three CAG, thirteen GIM, nine tumor-adjacent controls, and ten primary gastric

tumors. Clinical phenotypic information (specimen location and histology), cell counts, and sequencing information are available in **Supplementary Table 3**.

4) **Spatial validation cohort:** For the spatial validation steps (spatial transcriptomics and single-molecule fluorescent *in situ* hybridization studies, **smFISH**), we used formalin-fixed paraffin-embedded (**FFPE**) tissue specimens from five patients (three OLGIM II, one OLGIM III, and one GC) at Stanford Health Care. OLGIM samples were procured from the antrum region and the GC sample was procured from the *incisura angularis*. Blocks were sectioned into 10 $\mu$m thick tissue sections and placed onto Visium barcoded slide (10X Genomics). Slides were stained with hematoxylin and eosin **(H&E)** and imaged. The H&E-stained sections were manually annotated by a board-certified pathologist at the glandular level for regions of normal base, normal pit, metaplasia, dysplasia, and carcinoma. The spatial transcriptomics assay was run directly on these H&E-stained sections. For the smFISH assays, we utilized unstained sections immediately adjacent (within 10 $\mu$m) to the H&E slide. Phenotypic description of the specimens used for spatial validation are available in **Supplementary Table 4**.

##### Standard (bulk) RNA-seq of gastric tissue samples

The gastric tissue was processed using the TissueLyser LT compact bead mill (Qiagen, Venlo, Netherlands) per manufacturer's protocol using 5 mm stainless steel beads in 600 µL lysis buffer. AllPrep® DNA/RNA/miRNA Universal (Qiagen) spin column was used to bind the DNA, and RNA was extracted from the flowthrough was then used to extract the per manufacturer's protocol. RNA quality was assessed using Qubit RNA Broad Range Assay Kit and Qubit Fluorometer system (Thermo-Fisher Scientific, Massachusetts, USA). To assess the quality of the RNA, the samples were analyzed using the LabChip GX system (RNA Assay—Standard Sensitivity Perkin Elmer) as per the manufacturer's protocol. RNA samples with an RNA quality score (RIN value) greater than 4 were then used for direct DNA library generation using the KAPA mRNA HyperPrep Library Preparation Kit (Roche). TruSeq DNA UD Indexes (Illumina) were used for adapter ligation. The KAPA Library Quantification Kit (Roche) was used for library quantification. Library quality was assessed using an iSeq 100 system and the i1 Reagent v2 (Illumina). Sequencing was conducted on an Illumina Novaseq.

**Bulk RNA-seq analysis (GAPS cohort)**

We split the RNA-seq data into 2 independent sets: a discovery cohort comprising 88 paired biopsies from 46 individuals (46 antrum, 42 body; 22 high-risk OLGIM, 66 low-risk OLGIM) and a validation cohort comprising 115 individuals (215 paired patient biopsies: 115 antrum, 100 body; 22 high-risk OLGIM, 193 low-risk OLGIM).

*Differential expression analysis*

We used `filterByExpr` to filter out lowly-expressed genes followed by TMM normalization method from edgeR package in the R statistical programming environment.[9,10] Next, we performed unsupervised clustering through hierarchical clustering and principal components analysis on both the discovery and validation cohorts to confirm preferential grouping of samples into high- and low-risk OLGIM groups (**Supplementary Figure 1A-D**).

A schematic of the data analysis pipeline for the bulk RNAseq data is shown in **Supplementary Figure 2**. For both the discovery and validation data sets, we performed differential expression analysis in the same fashion. Specifically, we utilized a factorial design strategy to compare high- and low-risk lesions for each anatomic region (body and antrum). In addition, most samples had matched patient biopsies from both antrum and body: for the discovery cohort this involved 42/46 samples (91.3%) and for the validation cohort this involved

100/115 samples (87%). We used `voom` and `duplicateCorrelation` from limma[11] to estimate patient-specific weights for the regression fit, and a fold-change threshold of 1.25 was used to identify differentially expressed genes. To adjust for batch effects in the validation data set, comprised of 3 batches, we incorporated the sequencing batch as a co-variable for the regression models. Significance was set at 0.05 (FDR-adjusted P-values). In addition, we calculated the significance of the double variable (interaction term between the risk strata and anatomic region) over the differential expression profiles.

*Weighted gene co-expression network analysis (WGCNA)*

WGCNA was conducted using WGCNA R package[12]. The top 15% most variable genes (N = 5797) were used as input for WGCNA and normalized using the VST method in DESeq2 R package.[13] A soft threshold of 16 was selected for the generation of signed weighted networks, to achieve a scale-free topology model fit $R^2 > 0.8$ and a mean connectivity < 100. Informative gene modules were identified based on module-trait relationship and hierarchical clustering (**Supplementary Figure 4**).

*Common genes between DEA and WGCNA*

The differentially expressed genes were intersected with genes from informative modules from WGCNA to identify a set of genes both i) significantly upregulated and ii) co-expressed, common to the body and antrum of the stomach (**Supplementary Figure 4D**). A total of 314 genes met these criteria. We conducted hierarchical clustering using the scaled expression levels of these 314 genes with ComplexHeatmap[14] in R. We identified 5 gene clusters defining high-risk GIM. We selected a subset of 105 genes from a distinct cluster with the highest Z-score (C-5) for further validation in the held-out testing set (**Figure 2A**).

*Validation of the high-risk signature*

To validate the high-risk gene expression signature, we performed differential expression analysis as described above, in the held-out testing set. Differentially expressed genes common to the body and antrum of the stomach were intersected with common differentially expressed and co-expressed genes from the body and antrum in the discovery cohort. This yielded 100/105 (95.23%) overlap between the discovery and validation sets. Functional annotation of these 100 genes was performed through over-representation analysis, with the `enricher` function in clusterProfiler[15,16] R package. Gene sets were imported into R from Molecular Signature Database (MSigDB) using R package msigdbr.

**Spatial transcriptomics assay**

The Visium Spatial for FFPE Human Transcriptome Gene Expression Kit Gene Expression Kit (version 1) (10X Genomics) was used to prepare libraries according to the manufacturer's protocol. Briefly, 10 $\mu$m thick tissue sections from FFPE blocks were placed on a Visium Spatial Gene Expression slide, deparaffinized and stained with hematoxylin and eosin before coverslip removal and decrosslinking. Slides were imaged using a Keyence BZ-X microscope. Probe hybridization, ligation, release, extension, and library construction were performed as per protocol using 17 cycles for sample index PCR, and libraries were sequenced on an Illumina Novaseq 6000.

**Data analysis for spatial transcriptomic results**

The Space Ranger (10x Genomics) version 1.2.1 `mkfastq` command was used to generate Fastq files, and Space Ranger version 1.2.1 `count` was used with default parameters and alignment to GRCh38 to perform image alignment, tissue detection, barcode and UMI counting, and generation of feature-barcode matrix. Tissue regions were annotated by a pathologist (author JS) on a hematoxylin and eosin histology images from the Visium slide.

Raw counts were imported into the Seurat R package (version 4.3.0) and low-quality spots that detected <500 genes were removed. Genes detected in 3 or fewer spots were excluded, and counts were normalized with `SCTransform`. Spots within anatomically distinct tissue regions were labelled by unsupervised clustering refined by pathologist annotations. Spots were first clustered by `FindClusters` (resolution = 1.4) on the first 20 principal components and clusters marking tissue regions were annotated by pathologist annotations. Tissue regions were refined by transferring pathologist annotations to Loupe Browser (10x Genomics) version 6.1.0 to relabel any spots that did not match the pathologist annotations. Raw counts were summed from spots from each region of each patient sample to generate a pseudobulk count matrix, which was preprocessed using edgeR TMM normalization method in R. The limma-voom strategy was utilized to determine differentially expressed genes between metaplasia and spots mapping to normal gland base and pit. Gene signature scoring for each spot was performed with the `AddModuleScore` function in Seurat. To compare the gene signature module scores between metaplasia and normal gland pit or base regions. Normality of the data was assessed using Shapiro-Wilk test, followed by Bartlett test for homogeneity of variance and Welch T tests or Kruskal-Wallis followed by Dunn test for group comparisons.

**RNA-seq analysis (TCGA cohort)**

The differential expression analysis between tumor and adjacent non-tumor tissues from the TCGA RNA-seq dataset was performed to identify which of the spatially-resolved high-risk genes continued to be expressed at similar or increased patterns in cancerous tissue. We had multiple metrics and characteristics from these cancer specimens. RNA-seq data from 448 samples, including patient-matched tumor and non-tumor tissues, was downloaded and processed using `GDCquery`, `GDCdownload` and `GDCprepare` functions from TCGAbiolinks R package.[4] The functions `assay` and `colData` from R package SummarizedExperiment were used to extract raw count matrices and sample metadata. We subset the cohort to include only intestinal-type tumors and their patient-matched control tissues (n = 198; 180 tumor and 18 matched non-tumor samples) prior to filtering and normalization. Differential expression analysis was conducted using the limma-voom strategy. We used patient IDs as blocking variables for the regression models.

**Single-cell RNA sequencing**

Tissue biopsies were dissociated using a combination of enzymatic and mechanical dissociation with a gentleMACS Octo Dissociator (Miltenyi Biotec), and the resulting cells were cryofrozen using 10% DMSO in 90% FBS (ThermoFisher Scientific, Waltham, MA) in a CoolCell freezing container (Larkspur, CA) at -80 ºC for 24-72 hours followed by storage in liquid nitrogen. The cells were rapidly thawed in a bead bath at 37 ºC, washed twice in RPMI + 10% FBS, and filtered successively through 70 μm and 40 μm filters (Flowmi, Bel-Art SP Scienceware, Wayne, NJ), washed, filtered, and counted using 1:1 trypan blue dilution. Cells were concentrated between 500-1500 live cells/μl. The scRNA-seq libraries were generated using the Chromium Next GEM Single Cell 5' version 2 protocol, targeting 10,000 cells with 14 PCR cycles for cDNA and library amplification. The sequencing was performed on an Illumina NovaSeq 6000.

**Data processing of scRNA-seq**

We used Cell Ranger (10x Genomics) version 5.0.0 `mkfastq` command to generate Fastq files and Cell Ranger version 3.1.0 `count` with default parameters aligned to GRCh38 to generate a matrix of unique molecular identifier (UMI) counts per gene and associated cell barcodes. Seurat (version 4.0.1)[17,18] was used to construct Seurat objects from each sample. Quality control filters were applied to remove low-quality cells expressing fewer than 200 genes or greater than 30% mitochondrial genes, as well as doublets with UMI counts greater than 5000 or 8000. We removed genes that were detected in less than 3 cells. Data were normalized using `SCTransform` and first 20 principal components with a resolution of 0.6 or 0.8 were used for clustering. We removed computationally identified doublets from each dataset using DoubletFinder (version 2.0.3).[19] The 'pN' value was set to default value of 0.25 as the proportion of artificial doublets and the 'nExP' was set to expected doublet rate according to Chromium Single Cell 3' version 2 reagents kit user guide (10x Genomics). These parameters were used as input to the `doubletFinder_v3` function with number of principal components set to 20 to identify doublet cells. Individual Seurat objects were merged and normalized using `SCTransform`. The data sets were integrated using a soft variant of k-means clustering implemented in the Harmony algorithm (version 0.1.0)[20], using the `RunHarmony` function. This reduction was used in both `RunUMAP` and `FindNeighbors` functions for clustering. The first 20 principal components and a resolution of 0.6 were used for clustering. The data from the "RNA" assay were used for all further downstream analyses with other packages, gene-level visualization, or differential expression analysis. The data were normalized to the logarithmic scale and the effects of variation in sequencing depth were regressed out by including "nCount_RNA" as a parameter in the `ScaleData` function. Cell lineages were identified based on marker gene expression. The `Heatmap` function from ComplexHeatmap[14], `FeaturePlot`, `DimPlot`, and `VlnPlot` functions from Seurat were used for visualization. We performed a secondary clustering analysis of the epithelial lineage and epithelial stem cell with integration across samples using Harmony and a cluster resolution of 0.6.

SingleR algorithm[21] was used to determine cell identity by mapping the transcriptome of each individual cell to a reference single-cell atlas[22] of normal stomach and duodenum. Counts from the reference atlas were normalized to the logarithmic scale and used as a reference for automated annotation per cell using SingleR (version 1.14.1)[21]. Raw counts were used to annotate test datasets. Labels were predicted for each cell in the test dataset using the 'SingleR' function to calculate the Spearman correlation for marker genes for the reference dataset identified with Wilcoxon Rank Sum test. Following automated label assignment using this method, we confirmed results by examining marker gene expression. Cell labels for 43 cells identified as Tuft or Paneth cells were reannotated to Goblet cells based on marker gene expression in keeping with current recommended best practices.[23] In addition, epithelial cells from 9 tumor-normal pairs from seven GC patients[5] were used to evaluate gene signature activity using the `AddModuleScore` function in Seurat with default parameters.

**Single-molecule fluorescent *in situ* hybridization**

The manufacturer's protocol was followed to perform the RNAscope HiPlex12 Reagent Kit v2 (488, 550, 650) Assay (Advanced Cell Diagnostics Cat. No. 324419). Gastric tissue sections adjacent to those used for the Visium spatial transcriptomics assay were evaluated to identify twelve RNA target genes. The tissue sections mounted on slides were baked for 1 hour at 60℃ and deparaffinized in xylene and ethanol. Target retrieval was performed using a steamer at 99℃ followed by a protease treatment (Protease III for 30 min at 40℃). Probes for the twelve genes were hybridized for 2 hours at 40℃ and negative and positive control probes were run in parallel to assess sample RNA quality. A series of amplifiers were hybridized for signal amplification of single RNA transcripts for three target genes at a time which was visualized by hybridization with the first set of three cleavable fluorophores (T1-T3). Fluorophores corresponded to AF488, Dylight 550, and Dylight650. The sections were incubated with FFPE reagent for 30 min at room temperature to reduce autofluorescence and then counterstained with DAPI for 30 seconds. The sections were mounted, a cover slip was placed with ProLong Gold Antifade Mountant (Invitrogen, Cat. #P36930) and the section imaged using a Leica DMI 6000. After imaging, the coverslips were removed in a 4X SSC buffer (Sigma Aldrich, Cat.

#SRE0068), and the fluorophores were cleaved using the cleaving solution from the kit. The sections were hybridized with the next set of fluorophores (T4-T6), incubated with FFPE reagent, counterstained with DAPI, and reimaged. This process was repeated four times until all 12 target genes were imaged. Next, hematoxylin and eosin staining was conducted. Whole slide images were acquired using Aperio AT2 whole slide scanner (Leica Biosystems Inc., IL, USA).

*Fluorescence imaging*

Imaging was performed on a Leica DMI 6000 microscope with a 40X (NA 0.85) air objective. The microscope was controlled using the Lecia Application Suite X (LAS X) v3.6.0 software. The exposure and gain were qualitatively calibrated for each channel (DAPI, Alexa Fluor-488, Dylight 550, Dylight 650) and re-adjusted each round. To capture whole-slide images of the samples, we used the Tilescan function in LAS X to automatically acquire many image tiles using the motorized stage. The image tiles were acquired with 20% overlap to enable downstream stitching into whole-slide images. These images were exported from LAS X in .lif file format.

*Image preprocessing*

We inputted the .lif files from each sample into MCMICRO,[24] a microscopy image processing pipeline for multiplexed images. Within the MCMICRO pipeline, we first corrected for uneven illumination by applying the BaSiC algorithm.[25] Next, we use ASHLAR[26] to simultaneously stitch individual image tiles into whole-slide-images and register channels across multiple imaging rounds using the DAPI channel as a reference. Finally, for each sample we exported a multiplexed whole-slide OME-TIFF image containing all channels across all imaging rounds. We viewed these images with QuPath[27] v0.4.3 and exported snapshots to display in figures.

**Software and package versions**

R version 4.1.2

R packages: limma_3.50.3, edgeR_3.36.0, clusterProfiler_4.2.2, DESeq2_1.34.0, WGCNA_1.71, msigdbr_7.5.1, SummarizedExperiment_1.24.0, Seurat_4.1.1.

### Supplementary Figure 1

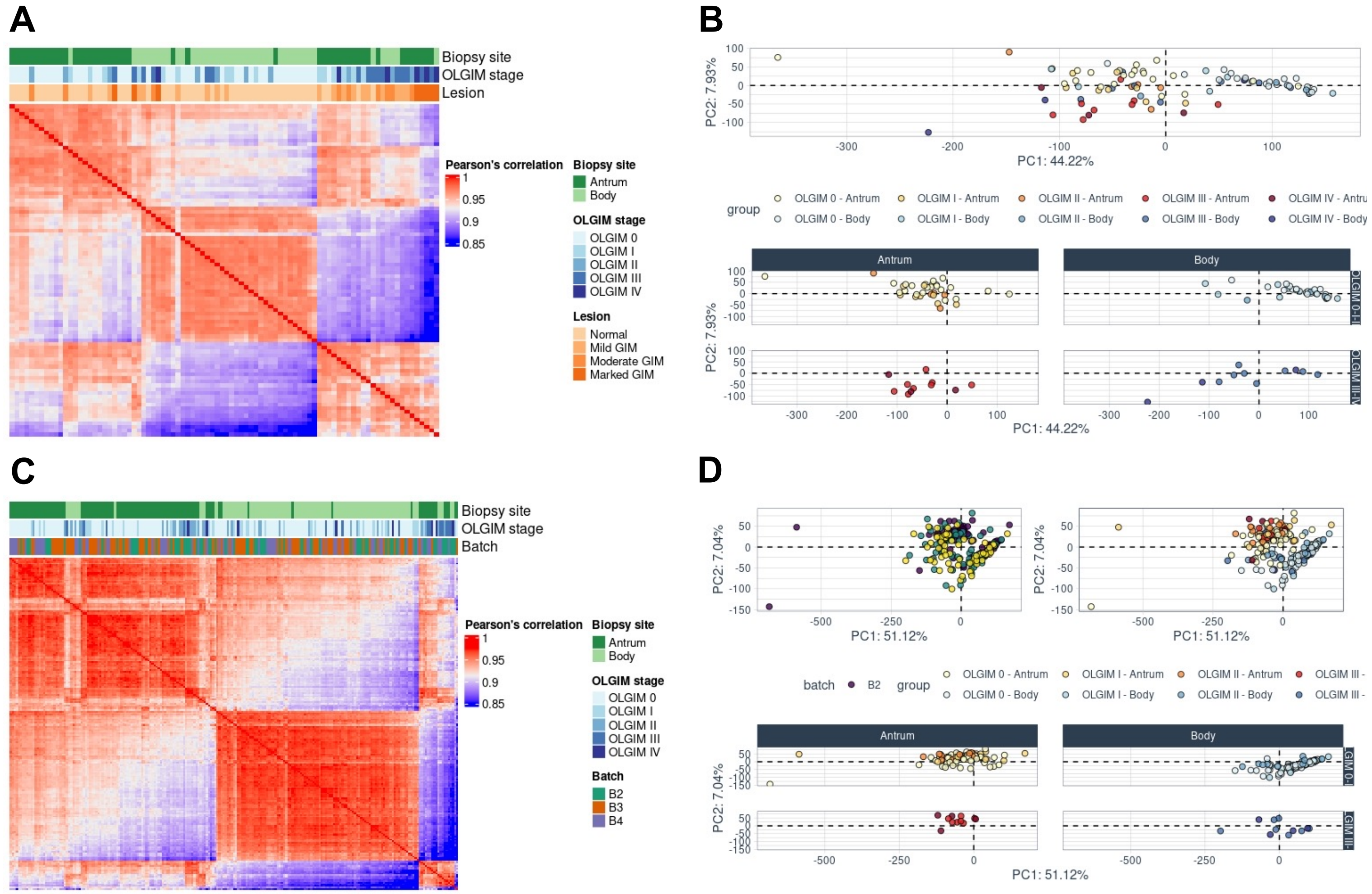


**Supplementary Figure 1. Unsupervised clustering.** A) Hierarchical clustering using pairwise Pearson's correlation coefficient between samples for the discovery cohort (N=88 samples: 22 high-risk, 66 low-risk). Complete linkage clustering method (default) was used. Preferential grouping of OLGIM III and IV samples is observed, regardless of anatomic site. OLGIM 0, I and II cases group preferentially by anatomic site of the biopsy (e.g., body or antrum). B) Principal components analysis showing preferential grouping OLGIM III and IV samples regardless of anatomic sites. OLGIM 0, I and II samples tend to cluster by anatomic site consistent with hierarchical clustering approach. C) and D) Hierarchical clustering and principal components analysis between samples for the held-out validation cohort (N=215 samples: 22 high-risk, 193 low-risk). Sequencing batch was added. No evident batch effects were observed by either method. B1: Batch 1, B2: Batch 2, B3: Batch 3.

### Supplementary Figure 2

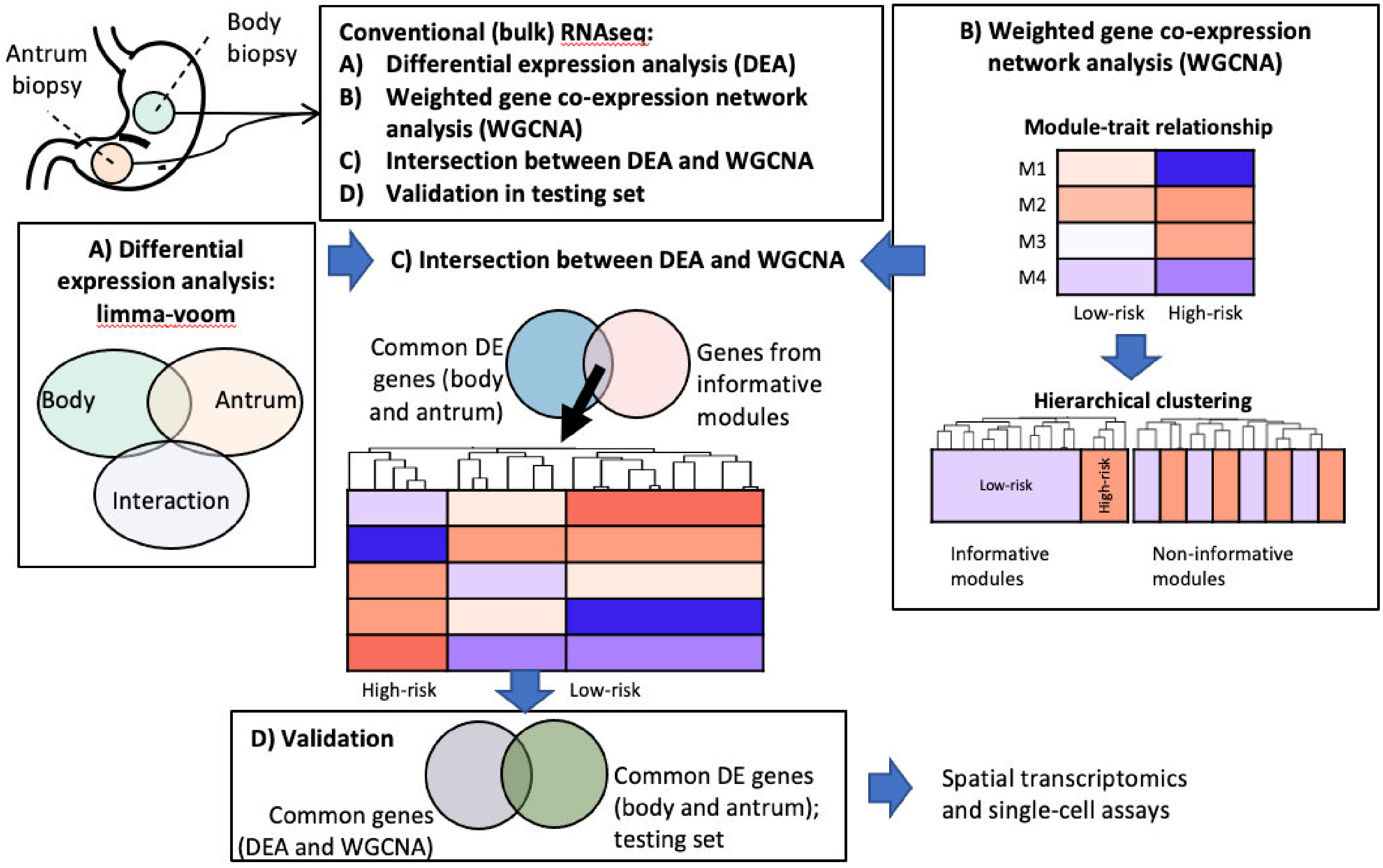


**Supplementary Figure 2. Analysis strategy for bulk RNA-seq data.** A) Differential expression analysis (discovery cohort). Samples grouped as high-risk (OLGIM III and IV) were compared with low-risk (OLGIM 0, I and II) in the antrum and body, separately. An interaction term was calculated by for genes which differential expression profile differed significantly by anatomic location (interaction term). Common differentially expressed genes in the body and antrum, excluding genes with a significant interaction term, were kept for downstream analysis. B) Weighted gene co-expression network analysis (WGCNA). WGCNA was conducted using top 15% most variable genes. Gene co-expression modules were identified. The first principal component of each gene module was calculated to compare high- and low-risk groups (module-trait relationship). Hierarchical clustering using scaled expression levels, Pearson correlation distance and Ward's clustering method was performed to inspect gene modules with module-trait relationship indicative of differences between high- and low-risk samples. Gene modules that clustered high-risk samples apart from low-risk samples were considered informative and kept for downstream analyses. C) Intersection between DEA and WGCNA. A total of 399 differentially expressed genes from A) were intersected with 815 genes from B). A curated set of 314 co-expressed and differentially expressed genes was identified. Based on gene clusters and Z-score, we selected a subset of 105 genes for further evaluation in the validation cohort. D) Differential expression was conducted as in A) in a held-out validation cohort of 215 samples. Differentially expressed genes from this validation cohort were intersected with the 105 genes from C). This resulted in the validation of a highly curated set of 100 genes (95.23% of the original genes).

#### Supplementary Figure 3

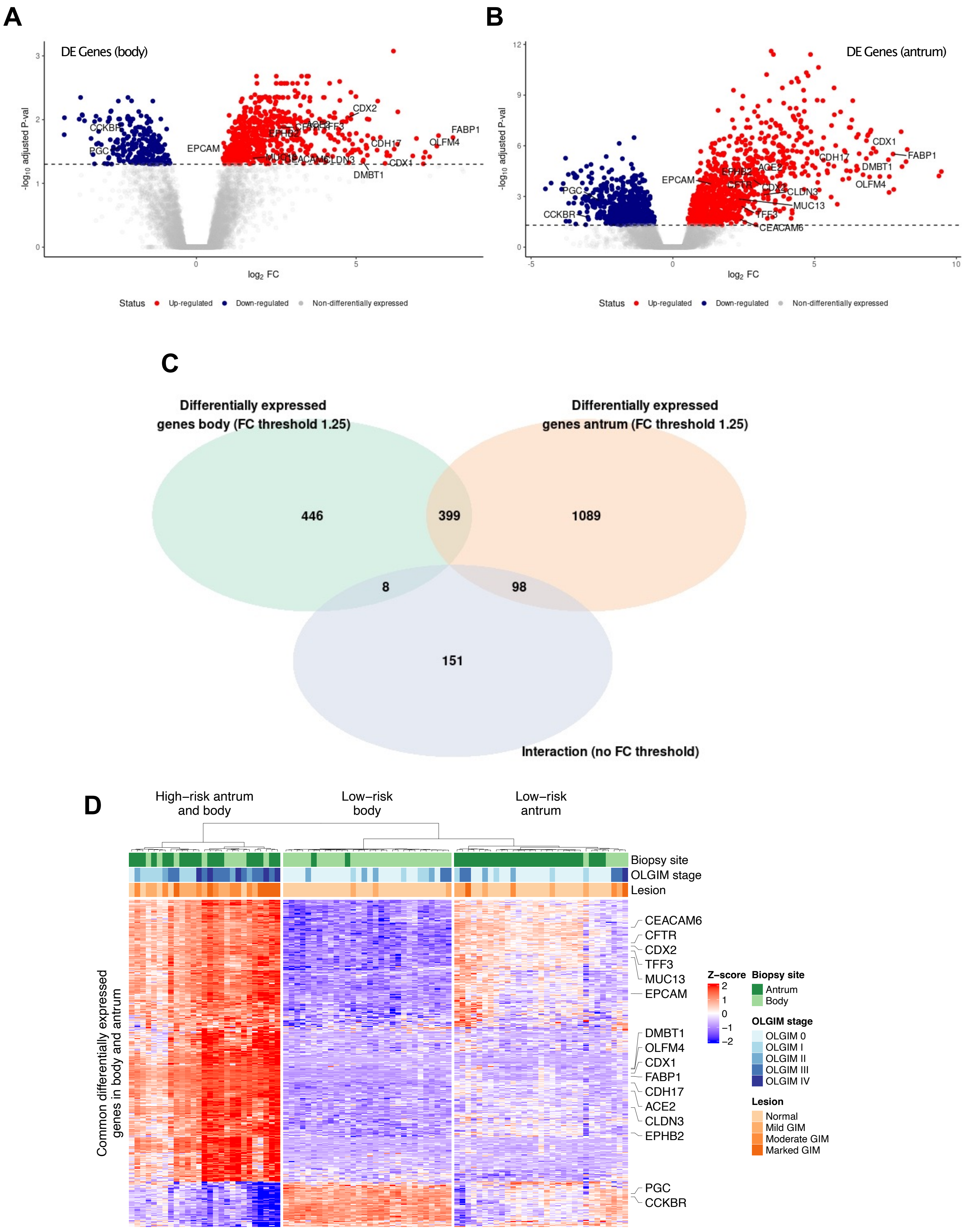

**Supplementary Figure 3. Differential expression analysis, discovery cohort.** A-B) Volcano plot showing significantly upregulated (red) and downregulated (blue) genes in the body (A) and antrum (B). C) Differentially expressed genes (DEGs) between high- and low-risk OLGIMs were defined using a fold-change threshold of 1.25 and adjusted p-value ≤ 0.05. DEGs for the body and antrum were identified. We further confirmed that this these intersected genes did not demonstrate any significant statistical interaction with anatomic location within the stomach. This analysis resulted in 399 genes. D) Heatmap and hierarchical clustering using Pearson distance and Ward clustering method of 399 differentially expressed genes common to the body and antrum. A-D: Known intestinal metaplasia and gastric epithelial markers are labelled.

### Supplementary Figure 4

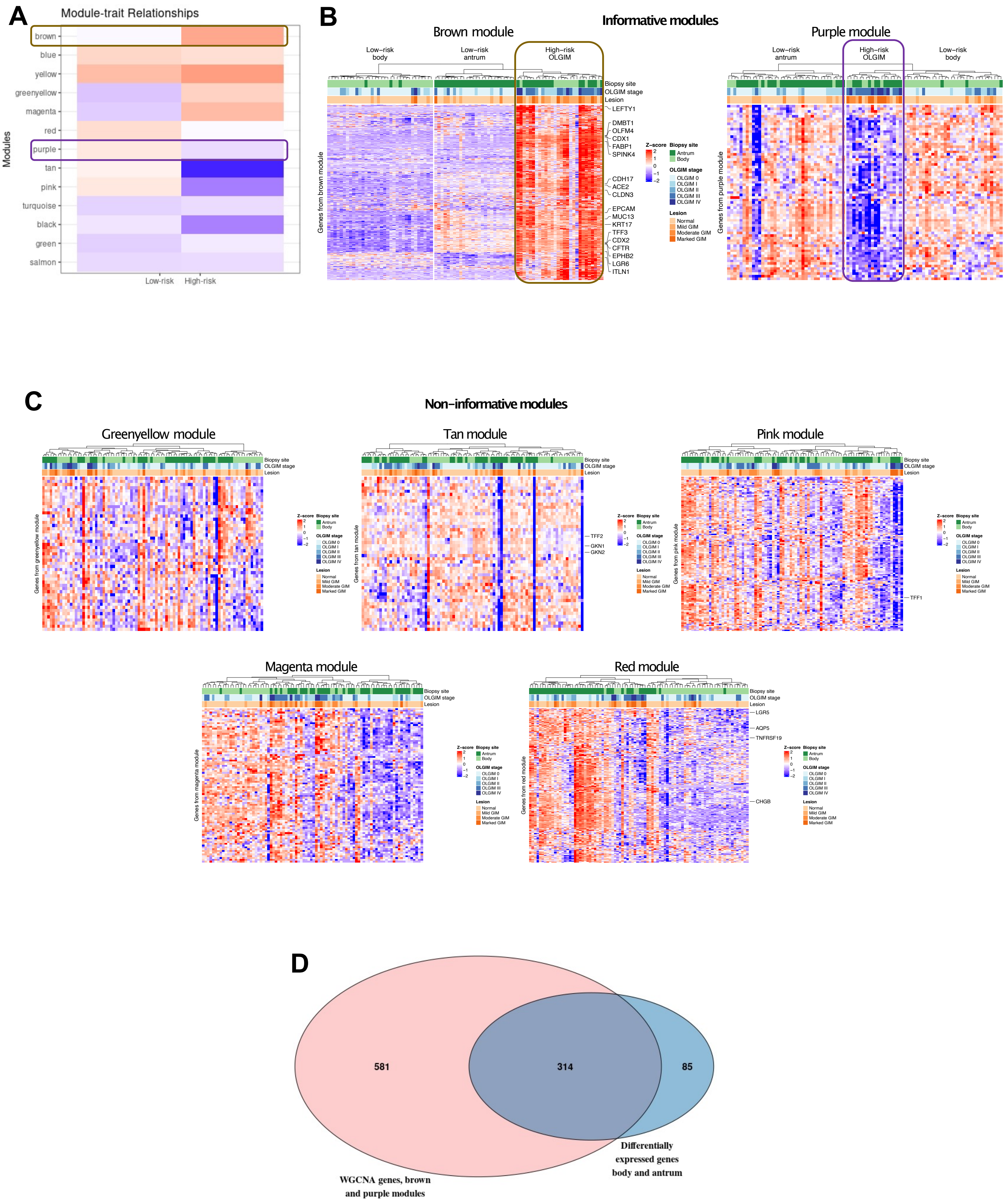

**Supplementary Figure 4. Weighted gene co-expression network analysis (WGCNA).** A) Module-trait relationship showing Z-scaled Eigen gene value (first principal component of each gene module) averaged across samples from high- and low-risk groups. Brown, green-yellow and magenta modules are increased in high-risk GIM. Red, purple, tan and pink modules are reduced in high-risk GIM. B-C) Hierarchical clustering of samples from brown and purple modules using Pearson distance and Ward clustering method. These modules show preferential clustering of high-risk GIM. The brown module captures most metaplasia and gastric epithelial cell markers, that are over-expressed in high-risk GIM. D-H) Non-informative gene modules from WGCNA. These modules do not segregate high-risk GIM. I) Intersection of genes from the brown and purple modules with results from differential expression analysis.

### Supplementary Figure 5

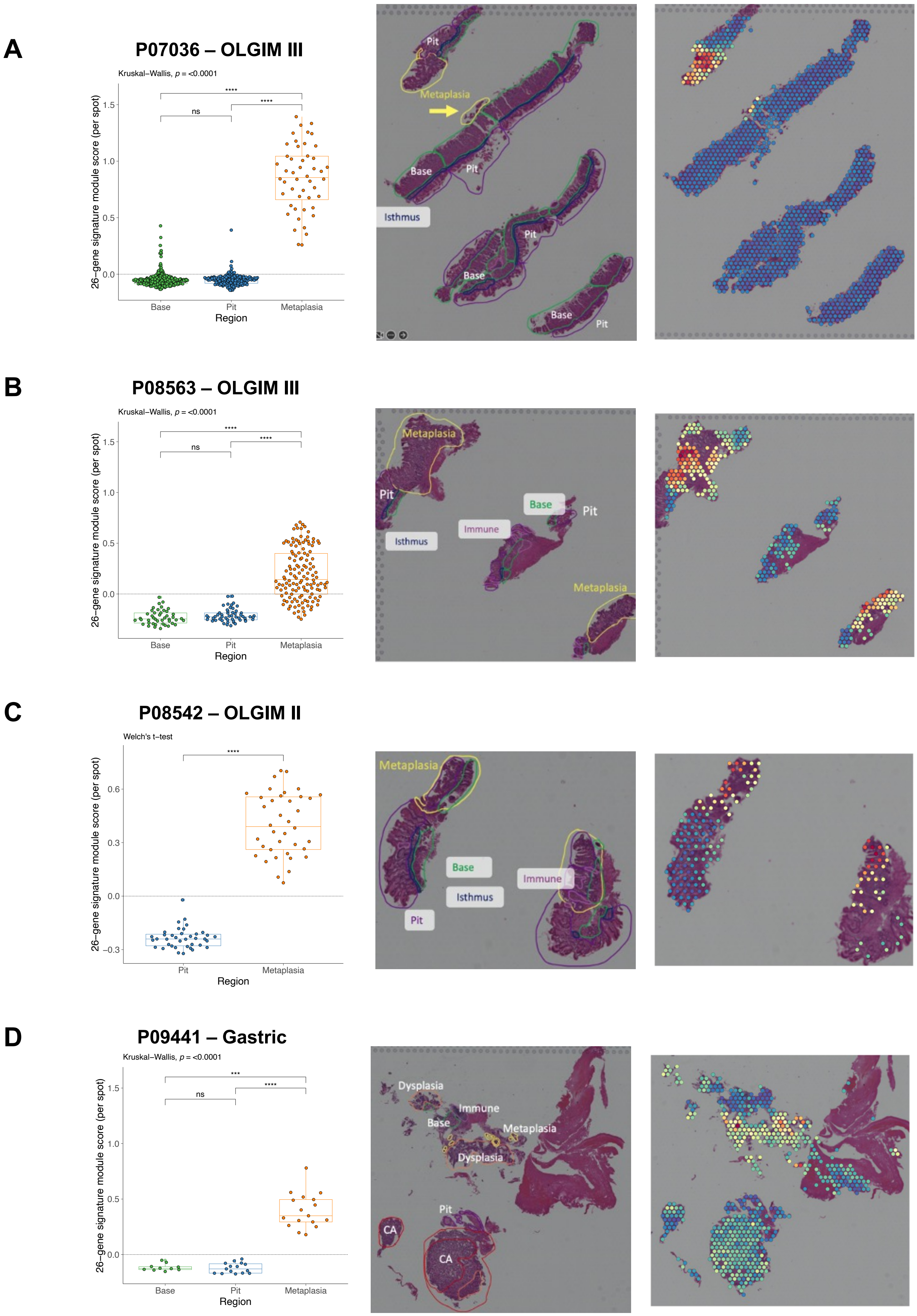

**Supplementary Figure 5. Spatial mapping of the module score.** The 26-module score value was calculated for each individual spot from 5 samples undergoing spatial transcriptomics assay. Statistically significant differences were observed between metaplastic foci and normal gland base and pit across all samples (Kruskal-Wallis followed by Dunn's test). Spatial mapping of the signature was highly consistent with pathologist annotations of intestinal metaplasia.

### Supplementary Figure 6

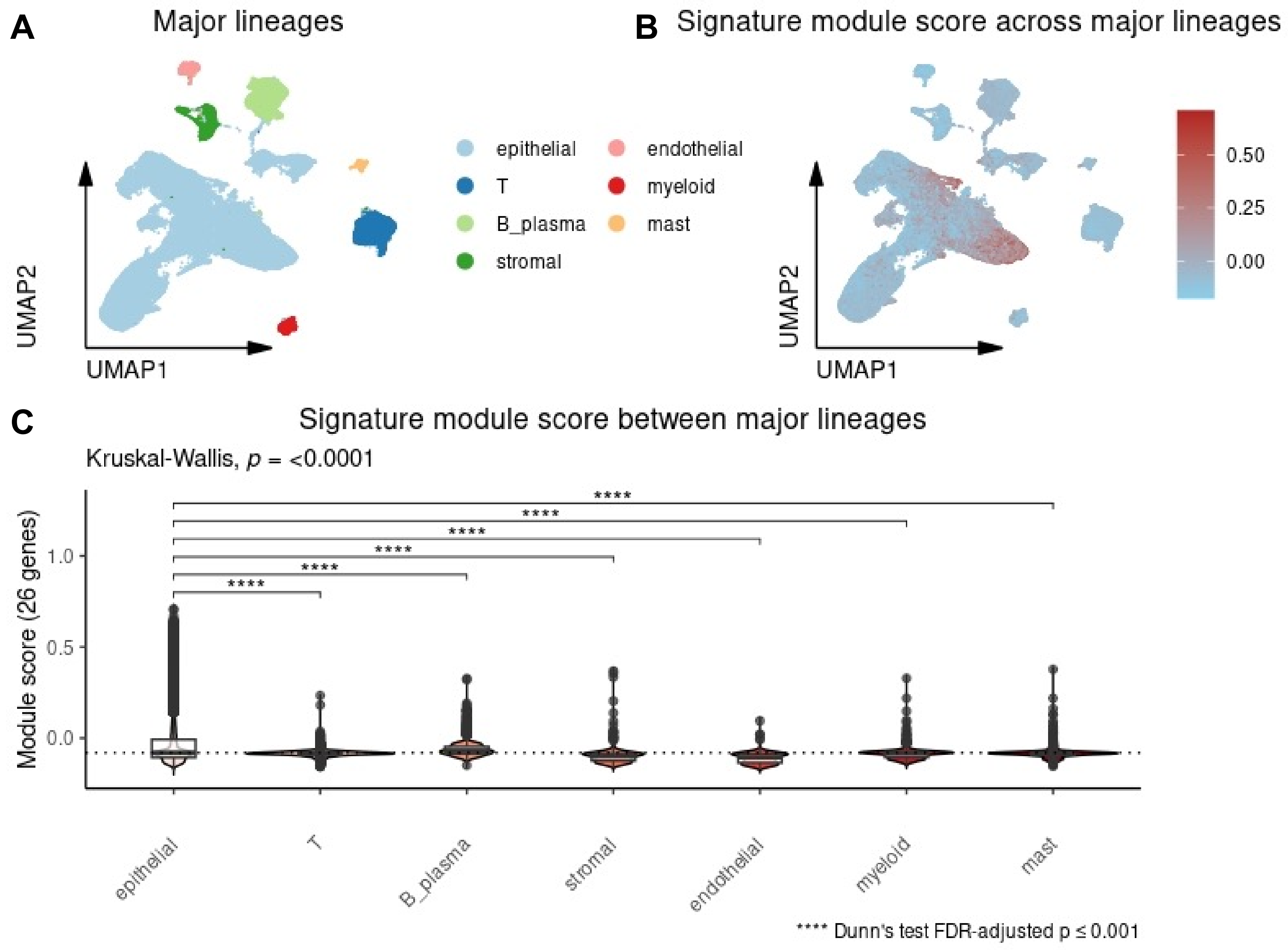


**Supplementary Figure 6. Signature module score is increased in epithelial cells.** A) UMAP plot showing 7 major cell lineages. B) Signature module score plotted across UMAP plot. C) Module score comparison between epithelial cells and other major cell lineages. Kruskal-Wallis followed by Dunn's test.

### Supplementary Figure 7

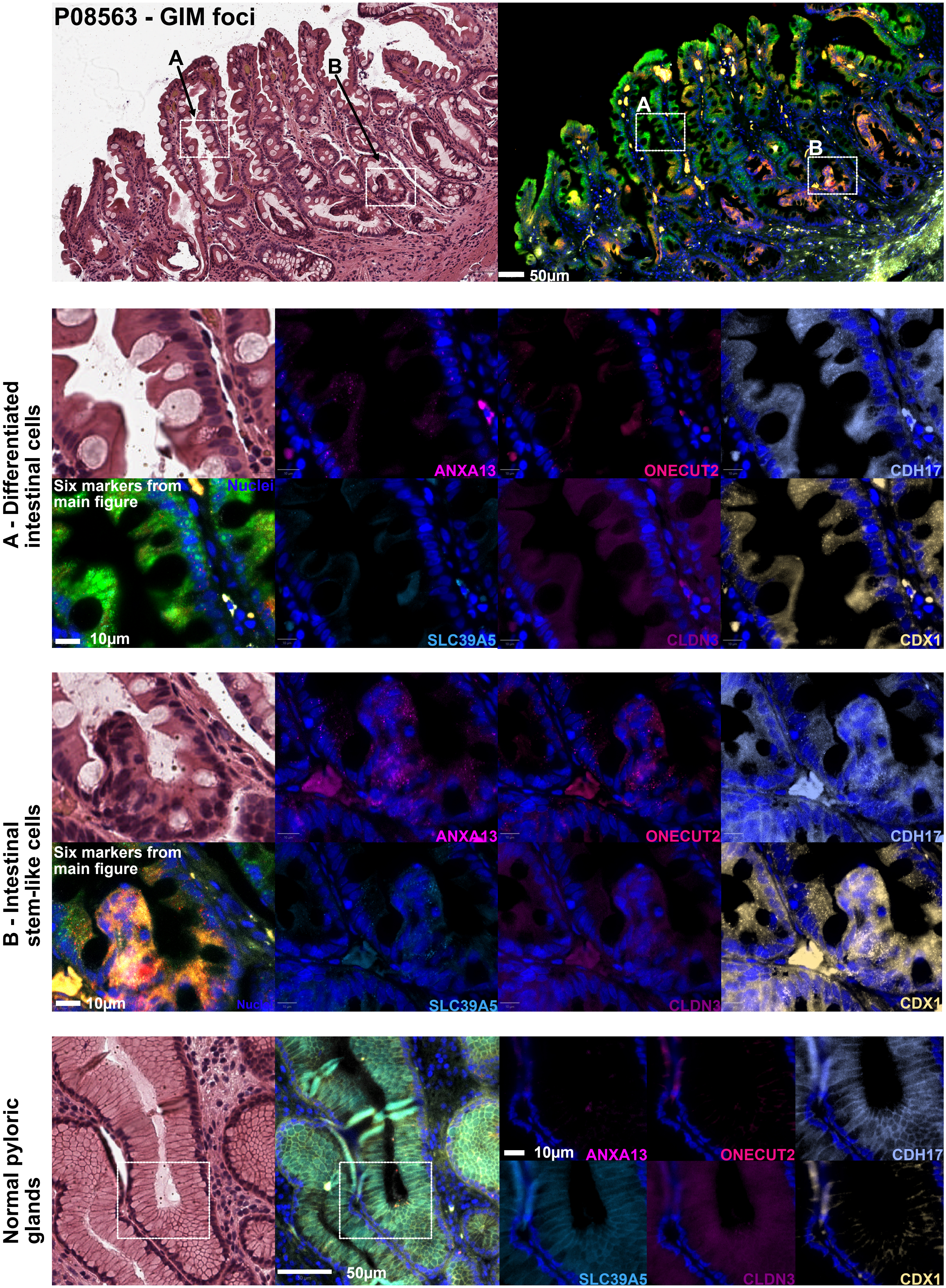

**Supplementary Figure 7.** H&E and corresponding RNAscope smFISH from patient P08563 (OLGIM III, antrum). Six genes from the signature are shown in the merged smFISH panels: TFF3, HKDC1, DMBT1, OLFM4, CPS1 and ANPEP. Regions A and B depict representative areas of well differentiated intestinal cells (A) and intestinal stem-like cells (B). Middle panels show inset magnification of regions A and B, with corresponding mRNA signals from the remaining 6 genes from the 12-gene signature: ANXA13, SLC39A5, ONECUT2, CLDN3, CDH17 and CDX1. The mRNA signals indicate that ANXA13, SLC39A5, ONECUT2, and CDX1 are expressed higher in stem-like cell foci than differentiated intestinal cells near the surface of the metaplastic glands. These mRNAs are not expressed in non-metaplastic normal glands from the same patient (bottom panels).

### Supplementary Figure 8

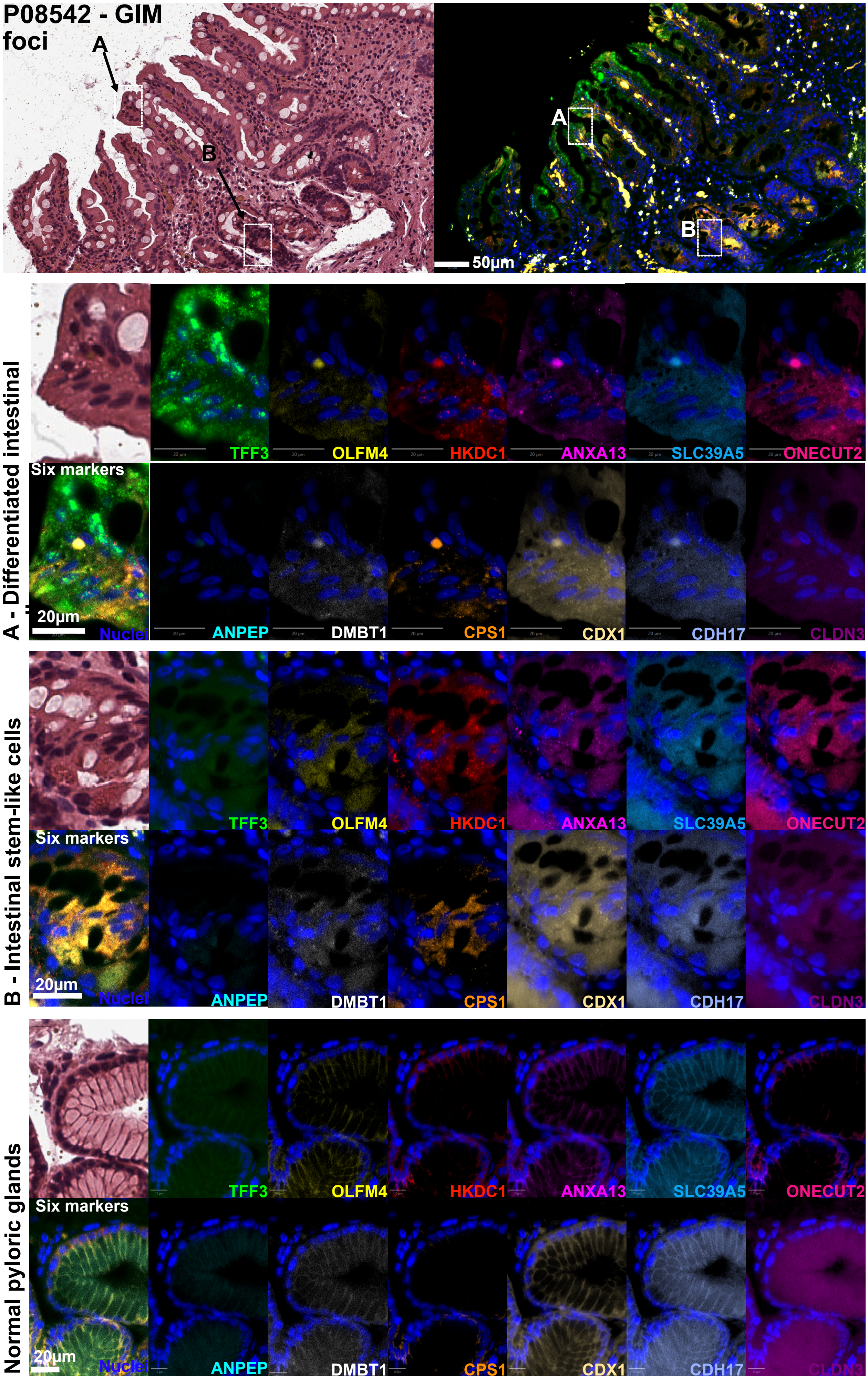

**Supplementary Figure 8.** H&E and corresponding RNAscope smFISH from patient P08542 (OLGIM II, antrum). Six genes from the signature are shown in the merged smFISH panels: TFF3, HKDC1, DMBT1, OLFM4, CPS1 and ANPEP. Regions A and B depict representative areas of well differentiated intestinal cells (A) and intestinal stem-like cells (B). Middle panels show inset magnification of regions A and B. The mRNA signals indicate that TFF3 is expressed mainly by differentiated intestinal lineages. OLFM4, DMBT1, HKDC1, CPS1, ANXA13, and CDX1 are expressed higher in stem-like cell foci than differentiated intestinal cells near the surface of the metaplastic glands. These mRNAs are not expressed in non-metaplastic normal glands from the same patient (bottom panels).

### Supplementary Figure 9

#### P07036 - Mixed foci

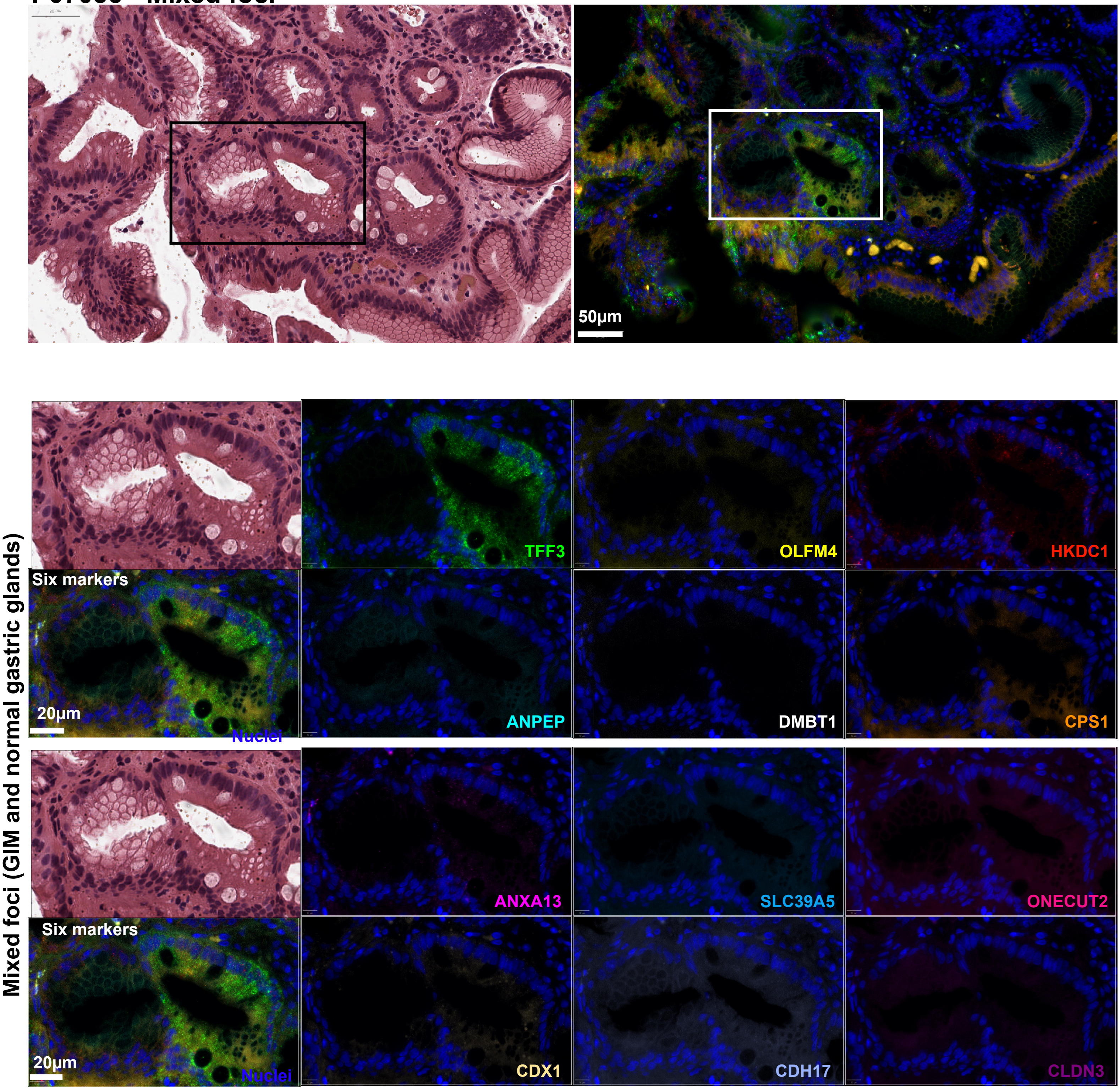

**Supplementary Figure 9.** H&E and corresponding RNAscope smFISH from a transverse section of metaplastic foci from patient P07036 (OLGIM II, antrum). Six genes from the signature are shown in the merged smFISH panels: TFF3, HKDC1, DMBT1, OLFM4, CPS1 and ANPEP. The highlighted region circumscribes immediately adjacent normal and metaplastic glands. The metaplasia-adjacent normal glands do not express any of the selected markers, whereas the metaplastic gland shows well-differentiated intestinal cells expressing TFF3 and, to a lesser amount, HKDC1, CPS1, ANXA13 and CDX1.

### Supplementary Figure 10

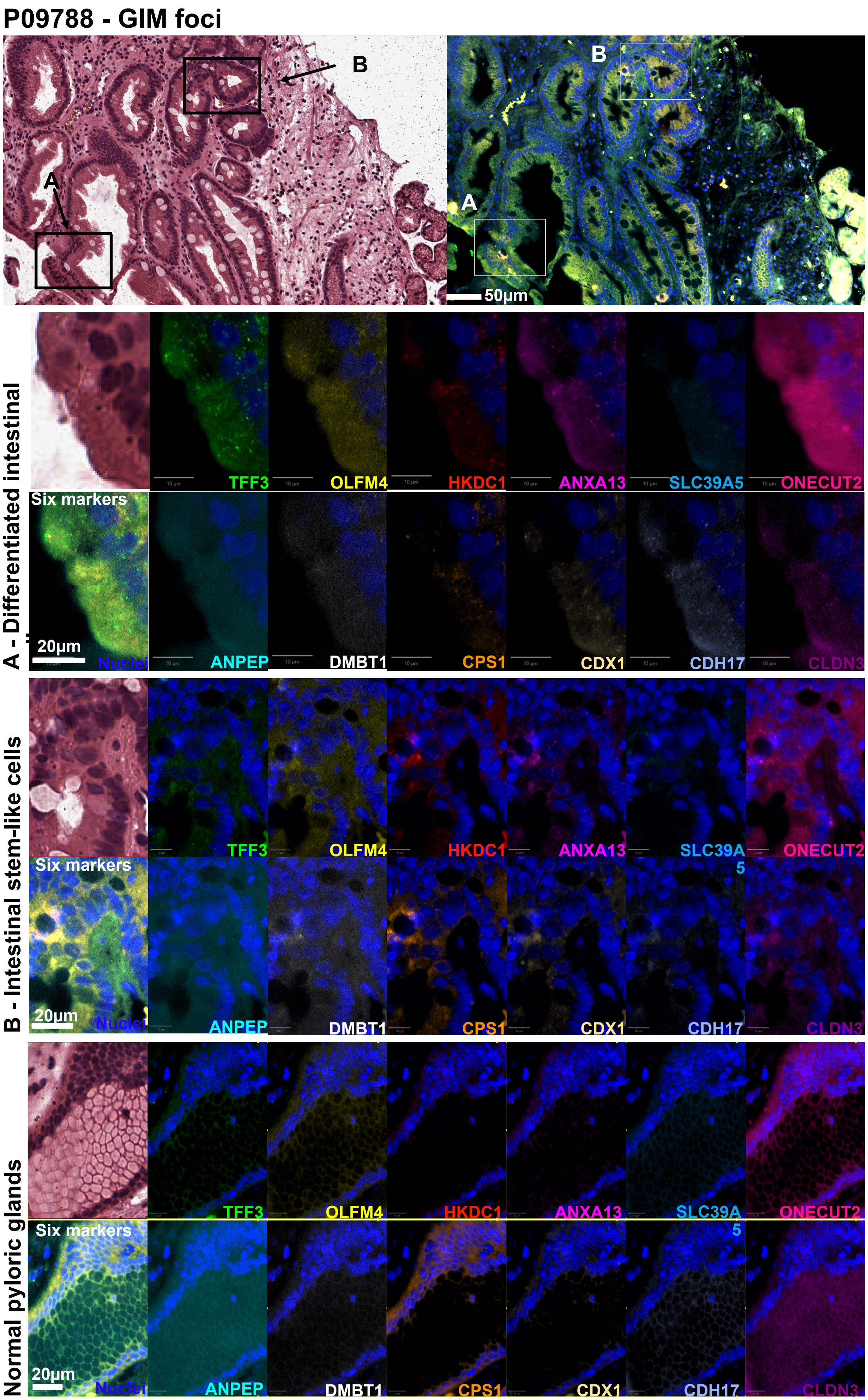

**Supplementary Figure 10**. H&E and corresponding RNAscope smFISH from an oblique section through metaplastic foci from patient P09788 (OLGIM II, antrum). Six genes from the signature are shown in the merged smFISH panels: TFF3, HKDC1, DMBT1, OLFM4, CPS1 and ANPEP. Regions A and B depict representative areas of well differentiated intestinal cells (A) and intestinal stem-like cells (B). Middle panels show inset magnification of regions A and B. The mRNA signals indicate that TFF3 is expressed mainly by differentiated intestinal lineages, that also show low expression of OLFM4, DMBT1, HKDC1, CPS1, ANXA13, CDX1, and CDH17. The stem cell-like region shows higher (moderate) expression of OLFM4, DMBT1, HKDC1, CPS1, CDX1 and ONECUT2 than differentiated intestinal cells near the surface of the metaplastic glands. These mRNAs are not expressed in non-metaplastic normal glands from the same patient (bottom panels).

### Supplementary Figure 11

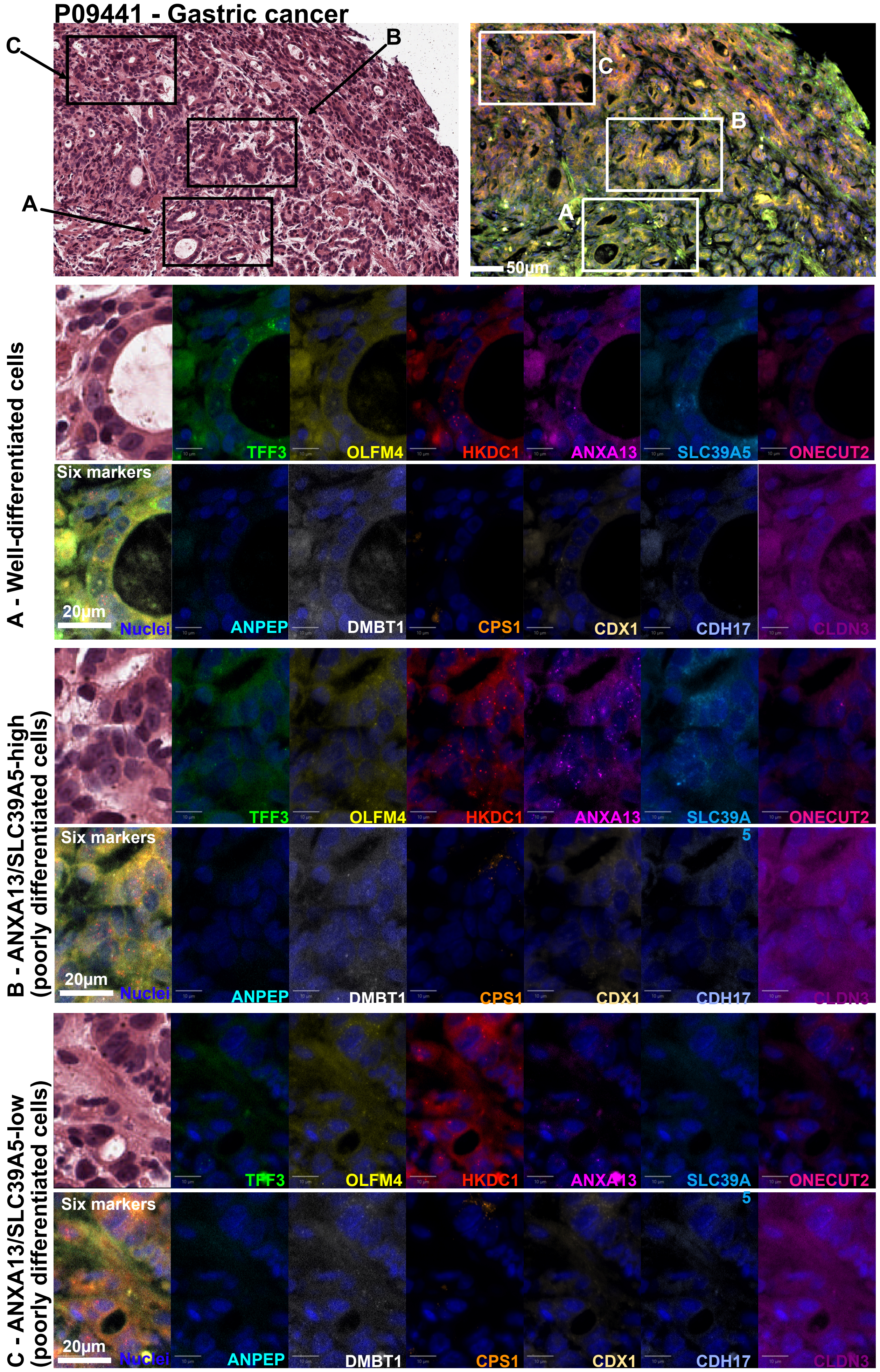

**Supplementary Figure 11.** H&E and corresponding RNAscope smFISH from patient P09441 (intestinal type gastric cancer, incisura angularis). Six genes from the signature are shown in the merged smFISH panels: TFF3, HKDC1, DMBT1, OLFM4, CPS1 and ANPEP. Three regions depict representative areas of well-differentiated tumor cells (A), poorly differentiated ANXA13/SLC39A5-high cells (B), and ANXA13/SLC39A5-low cells (C). smFISH images show co-expression of 12 high-risk gene transcripts in individual panels.
